# The beneficial effects of vaccination on the evolution of seasonal influenza

**DOI:** 10.1101/162545

**Authors:** Frank T. Wen, Anup Malani, Sarah Cobey

**Affiliations:** Department of Ecology and Evolution, University of Chicago, Chicago, Illinois, USA; The University of Chicago Law School, University of Chicago, Chicago, Illinois, USA; The University of Chicago Pritzker School of Medicine, University of Chicago, Chicago, Illinois, USA; Committee on Microbiology, University of Chicago, Chicago, Illinois, USA

## Abstract

Although vaccines against seasonal influenza are designed to protect against circulating strains, by affecting the emergence and transmission of antigenically divergent strains, they might also change the rate of antigenic evolution. Vaccination might slow antigenic evolution by increasing immunity, reducing the chance that even antigenically diverged strains can survive. Vaccination also reduces prevalence, decreasing the supply of potentially beneficial mutations and increasing the probability of stochastic extinction. But vaccination might accelerate antigenic evolution by increasing the transmission advantage of more antigenically diverged strains relative to less diverged strains (i.e., by positive selection). Such evolutionary effects could affect vaccination’s direct benefits to individuals and indirect benefits to the host population (i.e., the private and social benefits). To investigate these potential impacts, we simulated the dynamics of an influenza-like pathogen with seasonal vaccination. On average, more vaccination decreased the rate of viral antigenic evolution and the incidence of disease. Notably, this decrease was driven partly by a vaccine-induced decline in the rate of antigenic evolution. To understand how the evolutionary effects of vaccines might affect their social and private benefits, we fitted linear panel models to simulated data. By slowing evolution, vaccination increased the social benefit and decreased the private benefit. Thus, in the long term, vaccination’s potential social and private benefits may differ from current theory, which omits evolutionary effects. These results suggest that conventional seasonal vaccines against influenza, if protective against transmission and given to the appropriate populations, could further reduce disease burden by slowing antigenic evolution.

## 2 Introduction

As seasonal influenza evolves from year to year, antigenic differences between previously and currently circulating strains contribute to low vaccine efficacy [1–4] and a high incidence of influenza illness [2,5]. While vaccines are updated regularly to accommodate antigenic evolution, it is also theoretically possible for vaccination to affect antigenic evolution [6,7]. Vaccine-driven evolution or strain replacement has been observed in several pathogens, including avian influenza [8], Marek’s disease in poultry [9], and pneumococcus [10], among others [7]. Regional differences in the frequencies of influenza strains in humans suggest an influence of seasonal vaccination [11]. But traditional estimates of the public health benefits of influenza vaccines tend to focus on the benefits of vaccination in the current season and assume viral evolution is unchanged by the vaccine [12–16]. Accounting for the potential evolutionary impacts of vaccines, however, may alter projected assessments of their long-term value.

In theory, seasonal influenza vaccines might affect antigenic evolution in several ways [17–21]. First, by reducing the prevalence of infection, vaccination reduces viral population size and the rate at which antigenic escape mutants arise. Second, vaccination increases the amount of immunity in the population. By reducing transmission rates, this increased immunity could reduce the growth rate or invasion fitness of escape mutants, and thereby the rate of antigenic evolution (SI 1.1, Eq. S19, Fig. S1). Finally, stochastic extinction is more common in smaller populations (i.e., they are dominated by genetic drift), which should further reduce the strength of selection. These mechanisms underlie predictions that a hypothetical universal influenza vaccine, assumed to protect equally well against all circulating strains, should reduce the rate of antigenic evolution [19]. However, conventional influenza vaccines might *accelerate* antigenic evolution if the vaccine is less effective against strains that compete with vaccine-targeted strains, leading to strain replacement or vaccine escape [20,21], as seen in other pathogens [8–10,22].

Evolutionary effects could change the individual-level and population-level benefits of vaccination, which we refer to as the private and social benefits, respectively. Vaccination confers a private benefit to vaccinated individuals by directly reducing their risk of infection: on average, the seasonal influenza vaccine reduces the within-season rate of clinical laboratory-confirmed influenza infections in healthy adult recipients by 41% (95% CI 36-47%) [23]. Vaccination also confers a social benefit to the host population by reducing the burden of disease, although these effects are rarely measured. In the United States, vaccinating children reduces the risk of influenza infection in unvaccinated household contacts by 30-40% [24,25], in the local community by up to 5-82% [26], and in a metropolitan county by up to 59% [27]. In Japan, vaccinating children confers reduces mortality in the elderly by 17-51% [28]. The valuation of private and social benefits changes according to how much vaccination decreases the burden of disease. Increases in vaccination coverage have positive private and social benefits until the level required for herd immunity, at which point the disease risk (in a closed population) becomes zero, and there is no further benefit to vaccination [29]. Vaccine-induced evolution might also change the relative sizes of private and social benefits. For example, if vaccines slow antigenic evolution and thereby further decrease incidence, then their social benefit increases relative to the evolution-free case. However, as the social benefit decreases the risk of infection for the unvaccinated, their private benefit may fall commensurately. Additional incentives might then be necessary to compensate for less frequent voluntary vaccination [30, 31]. However, the private benefit may also increase as slower antigenic evolution improves the antigenic match between the vaccine and circulating strains.

Empirical estimates of the benefits of vaccination have so far been unable to measure the potential long-term evolutionary effects of vaccination. Most studies estimating the value of vaccination occur in temperate populations in North America, Europe, and Oceania, which have relatively high vaccine coverage but do not consistently contribute to influenza’s long-term evolution [32–36]. By contrast, source populations that contribute more to influenza’s evolution (e.g., China and India) have little vaccination [32–34] and few such studies of vaccination occur there [37].

We consider here the consequences of an idealized vaccination strategy in which vaccination occurs in populations that contribute to influenza’s long-term evolution. Such a scenario might arise if conventional seasonal vaccination becomes widespread in so-called “source” populations (e.g., East, South, and Southeast Asia [34]). To assess the potential effects of vaccination on antigenic evolution, we simulated the evolutionary and epidemiological dynamics of an influenzalike pathogen. We evaluated how different vaccination rates may slow antigenic evolution and in turn decrease the total burden of disease. We then quantified how the evolutionary effects change the relative magnitude of the private and social benefits of vaccination in the short and long term.

## 3 Methods

### 3.1 Modeling approach

To understand how vaccination on a global scale could affect influenza’s long-term evolution, we adapted an agent-based model to simulate the transmission and evolution of an influenza A/H3N2-like pathogen over 20 years in a well-mixed population [38]. We used a simple single-population model to test general principles in a hypothetical scenario where vaccination occurs globally.

In each time step of a tau-leaping algorithm, individuals can be born, can die, can become infected after contacting other hosts, can recover from infection, or can be vaccinated. Transmission occurs by mass action, with the force of infection given by

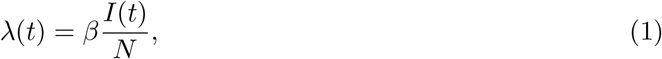

where *I* is the number of infected hosts. For computational efficiency, individuals cannot be coinfected.

Antigenic phenotypes are represented as points in 2-dimensional Euclidean space (Fig. 1A). This space is analogous to the main components after multidimensional scaling of pairwise measurements of cross-reactivity in hemagglutination inhibition (HI) assays, where one antigenic unit of distance represents a twofold dilution of antiserum [39,40]. One antigenic unit corresponds to a two-fold antiserum dilution in a hemagglutination inhibition (HI) assay. At the beginning of the simulation, a single founding strain is introduced at the endemic equilibrium in the host population. When hosts recover from infection, they acquire lifelong immunity to the infecting strain. In reality, immunity to infecting strains appears to last on the order of decades, if not longer [41–43]. Upon contact with an infected host, the probability that the susceptible host becomes infected is proportional to the distance *d*_n_ between the infecting strain and the nearest strain in the susceptible host’s infection history, with one unit of antigenic distance conferring a 7% absolute increase in risk (Eq. 3) [1,38,44].

**Figure 1:**
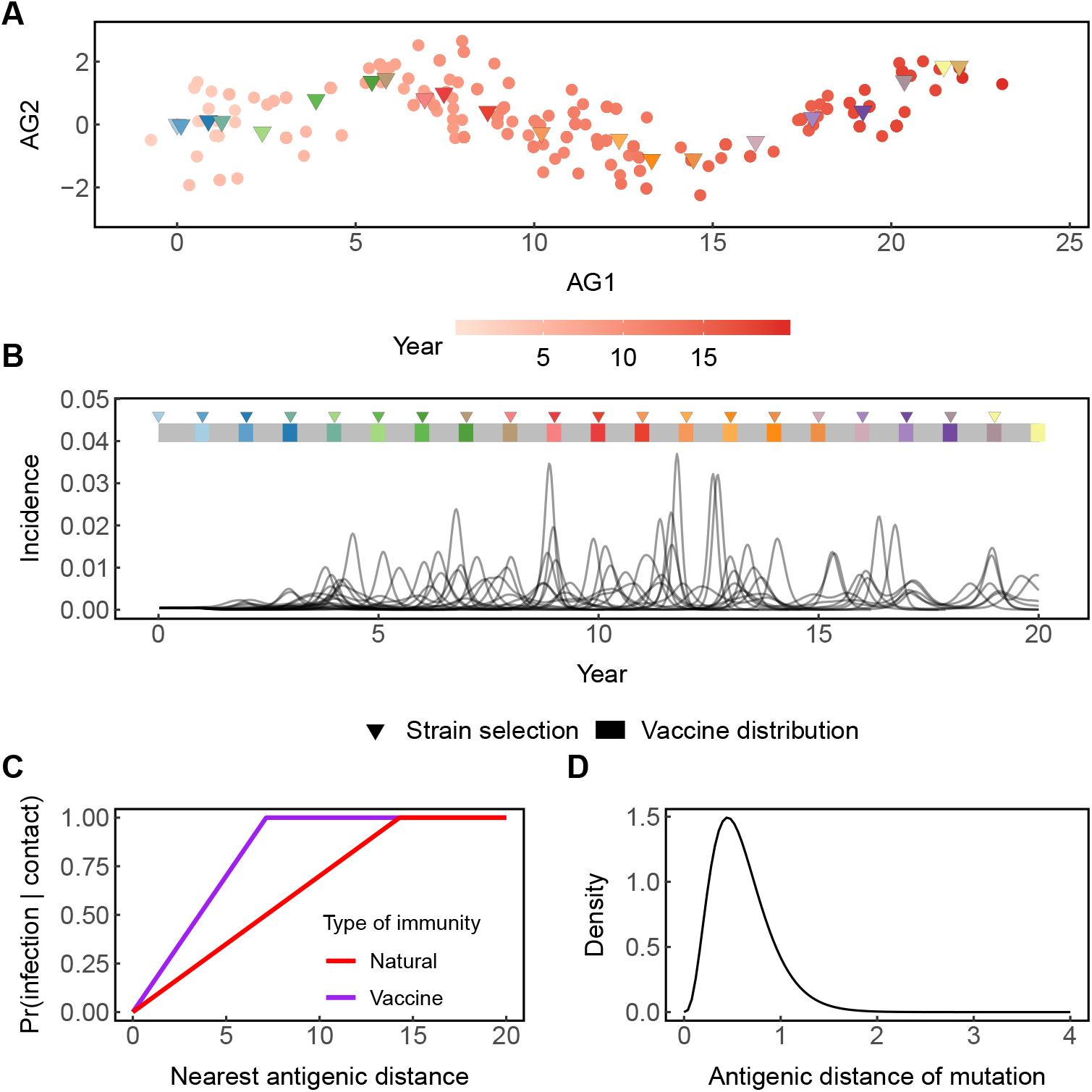
Properties of the model. (A) Antigenic phenotypes are represented as red points in two-dimensional space (AG1 is antigenic dimension 1 and AG2 is antigenic dimension 2). The shading of the points corresponds to the time that the strains appear. Over time, new strains appear as old strains can no longer transmit to immune hosts. Viral evolution is mostly linear in antigenic space. The amount of evolution is calculated as the distance between the founding strain and the average phenotype of strains circulating at the end of the simulation. Vaccine strains (triangles) are chosen at the beginning of each year by averaging the antigenic phenotype of all circulating strains. (B) Incidence per 10 days for 20 replicate simulations. Cumulative incidence (not shown) is calculated as the sum of cases over the duration of the simulation. Although the depicted model output is without vaccination, a hypothetical vaccine distribution schedule is shown by the bars and triangles. Strain selection occurs on the first day of each year. The vaccine is then distributed beginning 300 days after strain selection for 120 days. The triangles indicate the time points of vaccine strain selection, and the matching colored bars indicate the corresponding window of vaccine distribution for the selected vaccine strain. (C) Upon contact, the risk of infection increases linearly with the distance between the infecting strain and the strain in the host’s infection or vaccination history that minimizes the risk of infection (Eq. 3). In this example, for illustrative purposes, vaccines confer half the breadth of natural immunity (*b* = 0.5). However, by default, we simulate vaccines that have the same breadth as natural immunity (*b* = 1.0). (D) The sizes of antigenic mutations are chosen from a gamma distribution. The radial directions (not pictured) of mutations are chosen from a random uniform distribution.

Each infection mutates to a new antigenic phenotype at a rate *μ* mutations per day. The mutation’s radial direction is drawn from a uniform distribution, and the size (distance) is drawn from a gamma distribution with mean *δ*_mean_ and standard deviation *δ*_sd_ (Table S1, Fig. 1D).

### 3.2 Model validation and choice of parameters

The model reproduces characteristic epidemiological and evolutionary patterns of the seasonal A/H3N2 subtype without vaccination (Fig. 1A,B). We investigated the credibility of the model *without* vaccination because the evolution of H3N2 appears driven by populations with negligible vaccination rates: the dominant source populations have nearly 0% vaccine coverage [32,34]. We chose transmission and mutation parameters (Table S1) such that simulated epidemiological and evolutionary patterns most resembled qualitative patterns and quantitative metrics observed for H3N2 (Table 1) [45]. H3N2 has remained endemic in the human population since its emergence in 1968 and also has low standing genetic and antigenic diversity. Due to the stochastic nature of the simulations, the viral population goes extinct 18% of the time and becomes too diverse 29% of the time across replicate simulations. A viral population is considered too diverse when the time separating two co-circulating lineages (time to most recent common ancestor, or TMRCA) exceeds 10 years [38,45], since recent H3N2 HA lineages have coexisted for no more than 7 years. The remaining 53% of simulations show qualitatively influenza-like dynamics that reproduce key epidemiological and evolutionary statistics of H3N2 (Table 1).

**Table 1:** Agreement between simulated and empirically measured epidemiological and evolutionary metrics of H3N2. Simulated values are averages over 20 replicate simulations.

| Metric | Simulated value | Empirical estimate |
| --- | --- | --- |
| TMRCA (years) | 3.80 (SD = 0.52) | 3.84 [34] |
| Antigenic evolutionary rate (antigenic units/year) | 1.09 (SD = 0.14) | 1.01 [40] |
| Annual incidence per person | 9.0% (SD = 1.0%) | 9-15% [46] |
| Time between infections (years, 1/annual incidence) | 11.1 (SD = 1.3) | 5-11 [47–49] |

That 47% of simulations are not H3N2-like does not necessarily imply that the model is inaccurate: the H3N2 lineage circulating since 1968 represents only a single instance of that subtype’s global evolution. Moreover, two lineages of influenza B emerged approximately 30 years ago and have co-circulated since, demonstrating an instance of high diversity in influenza. The unusually high diversity of currently co-circulating H3N2 lineages suggests it might be capable of similar dynamics, which were not foreshadowed by the prior few decades of observation [50]. We also find agreement between the model’s epidemiological dynamics when comparing against analytic expectations without evolution (analytic solutions for a model with evolution are intractable), which indicates that the transmission dynamics behave as expected (Supplement 1.2, Figs. S2, S3, S4, and S5). The extinctions are attributable to stochastic amplification of epidemics, which is a common feature of nonlinear models [51]. A metapopulation structure might provide some buffer against extinctions [52], but would not change the effects of vaccination, assuming that vaccination is distributed evenly in space. We therefore implement a simple population structure to make general predictions about vaccination on a global scale.

### 3.3 Modeling vaccination

To assess the potential effects of vaccination on antigenic evolution and disease burden, we introduced vaccination to the host population. Vaccination occurs at rate *r*, breadth *b* (relative to natural immunity), and lag *θ* (relative to the timing of strain selection). The vaccine strain is selected on the first day of each year. The antigenic phenotype of the vaccine strain is the average (in 2D antigenic space) of contemporaneous circulating strains. In reality, strains are considered for inclusion in the vaccine if they are considered likely to spread (e.g those that circulate at high frequency or those that are highly antigenically diverged) [53]. By default, the vaccine is distributed for 120 days. This schedule approximates vaccine distribution in the United States, which usually runs from September through February and peaks in October or November, 8-9 months after strain selection [53]. During the period of vaccine distribution, individuals are randomly vaccinated at a constant daily rate according to the specified annual vaccination rate.

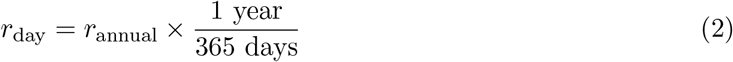

Vaccine recipients are selected at random with replacement, so approximately 4.9% of individuals in the population are vaccinated every year at a 5% annual vaccination rate. Since individuals are randomly vaccinated each year, the fraction of ever-vaccinated individuals increases from one season to the next. At a 5% annual vaccination rate, ~48.4% of the population has been vaccinated at least once by the twentieth year (Fig. S6A). At this rate, vaccination effectively renders 26.0% of individuals immune when vaccination is in equilibrium with antigenic evolution (Fig. S6B).

We also tested the effects of the breadth of immunity conferred by vaccination. The vaccine’s breadth *b* is defined as the ratio of the vaccine-induced immunity to that of infection-induced (or “natural”) immunity (Fig. 1). Vaccines with *b* = 1 have breadth identical to natural immunity, whereas vaccines with *b* < 1 (*b* > 1) have respectively smaller (larger) breadth compared to natural immunity. Thus, a host’s probability of infection upon contact is given by

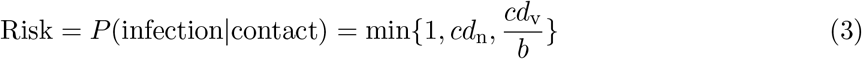

where *d*_n_ is the distance between the infecting strain and the nearest strain in the host’s infection history, and *d*_v_ is the distance between the infecting strain and the nearest strain in the host’s vaccination history (if the host is vaccinated) and *c* = 0.07 is a constant for converting antigenic distance to a risk of infection, derived from vaccination studies [1, 38, 44]. By default, the breadth of vaccine-induced and natural immunity are equal (*b* = 1).

### 3.4 Model output

We quantified vaccination’s effects on viral evolution and epidemiology using four metrics: cumulative antigenic distance evolved, cumulative incidence, the probability of excessive diversity, and the probability of extinction. First, because influenza evolves roughly linearly in two antigenic dimensions [38–40], we measured the cumulative amount of antigenic evolution by calculating the antigenic distance between the founding strain’s antigenic phenotype and the average antigenic phenotype of strains circulating at the end of the simulation (Fig. 1). We only estimated cumulative antigenic evolution in simulations that were not too diverse, since this metric is inadequate for viral populations with deep branching. Second, we measured the burden of disease by calculating the cumulative incidence, or the total number of cases over the duration of the simulation divided by the population size (Fig. 1). Third, we calculated the probability that viral populations would become too diverse (TMRCA > 10 years), since vaccination may qualitatively alter evolutionary patterns. Viral populations that are too diverse can cause high incidence because hosts are unlikely to have immunity against distant antigenic variants. Fourth, we calculated the probability of extinction by calculating the fraction of simulations that went extinct out of 500 replicates.

### 3.5 Measuring evolutionary effects of vaccination

To estimate the contribution of evolution to vaccination’s epidemiological impact, we compared simulations in which vaccination could affect antigenic evolution to simulations where it could not (Fig. S10). To generate the latter, we created a simulation where vaccination could not affect antigenic evolution, the “static” simulation (Fig. S10). We first ran 500 simulations of the model without vaccination to be used as a reference. For each simulation, we recorded the circulating strains and their relative abundances at each time step to use as reference viral populations. The evolution of these reference viral populations is unaffected by vaccination since they were obtained from simulations without vaccination.

To run the static simulation where vaccination could not affect antigenic evolution, we first randomly selected one of the reference viral populations. In each time step of the static simulation, the composition of the viral population was replaced with that of the reference viral population at the matched time step, scaled for prevalence. In this way, vaccination could still alter the overall viral abundance, but the rate of antigenic evolution had already been set by the dynamics of the simulation without vaccination. Thus, vaccination was separated from the evolutionary process.

### 3.6 Estimating the private and social benefits of vaccination

A linear panel regression model was fitted to simulated panel data to identify the private and social benefits of vaccines over 20 years. The social benefit is also known as the “indirect effect” and the private benefit is also known as the “direct effect” as defined in [54].

To generate panel data, we ran simulations at six annual vaccination rates *r* (0%, 1%, 3%, 5%, 7%, and 10%) and recorded individual hosts’ dates of infection and vaccination. We ran 20 replicates for each unique combination of rate and breadth, and randomly sampled 2,500 individuals (0.005% of the entire host population) at the end of each simulation for analysis, yielding up to 2 million observations for each fitting. If a simulation was terminated early because the virus went extinct before 20 years, additional data points were filled in according to the initial vaccination rate and assuming no new infections. We fitted a linear panel model (equation 4) to the simulated longitudinal vaccination data from multiple simulations *j*. Observations are at host *i* level in each season *τ* (see Table S2 for hypothetical example). The dependent indicator variable *V_ijρ_* equals 1 if a host is infected at any point in the current season *τ* and 0 otherwise. The indicator *V_ijρ_* equals 1 if a host is vaccinated in the current season. Analogously lags *V_ijρ-k_* measure vaccination in period *τ– k*. The vaccination rate indicators *R_rij_* equal 1 if the annual vaccination in the host population is equal to *r*% (e.g., when the rate is 5%, then *R*_5*ij*_ = 1). The regression is estimated as a linear panel model (with random effects) in order to simplify interpretation of reported coefficients. Standard errors are clustered at the simulation-level to account for correlation in outcomes across hosts in a simulation. The estimated equation is:

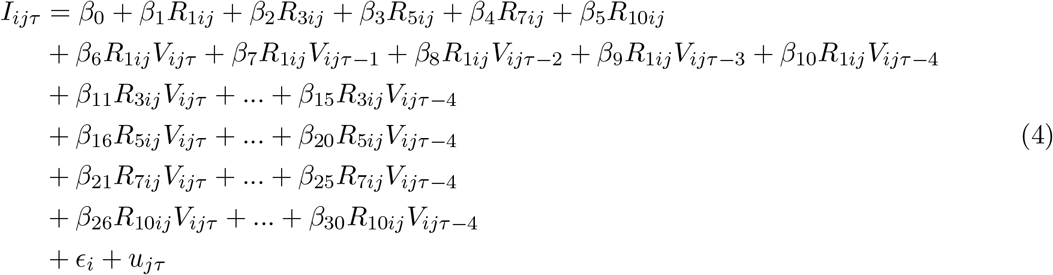

The fitted coefficients estimate the absolute change in the probability of infection given the host population’s vaccination rate and an individual’s vaccination status. We converted these absolute risks to odds ratios in keeping with standard reporting of influenza vaccine effectiveness. For example, 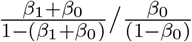 gives the odds ratio of infection for an *unvaccinated* individual’s risk of infection in the current season when the population vaccination rate is 1% relative to the odds of infection in an unvaccinated population. The same formula applied to *β_x_* for *x* ∈ {1,2, 3, 4, 5} represents the social or indirect benefits of vaccination under different vaccination policies.

The model is interacted (*β*_4_ to *β*_30_) to estimate the private benefit for each vaccination rate. Thus, 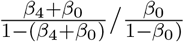 gives the ratio of the odds of becoming infected in the current season for a host who has been vaccinated in the current season and is in a population with an annual vaccination rate of 1% relative to the odds of infection for a host who is in a population with a 1% vaccination rate but has not been vaccinated in 5 years. Likewise, 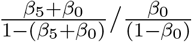 estimates the ratio of odds of becoming infected in the current season given vaccination one season ago and living under a 1% vaccination rate policy relative to an unvaccinated host also living under a 1% vaccination rate policy. More formally, 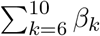 is the impulse response to vaccination over 5 years and measures the total individual-level protective benefit of vaccination over time when the vaccination rate is 1%. The same reasoning applies to the terms associated with the other two vaccination rates.

We also estimate the benefits of vaccination directly from incidence to validate the regression model. To estimate the social benefit (the indirect effect in [54]) for a specific vaccination rate r, we calculate the ratio of the odds infection for an unvaccinated host in a vaccinated population relative to the odds of infection in an unvaccinated population. For *x* ∈ {1, 2, 3},

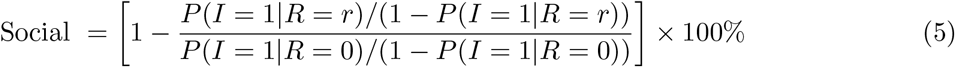

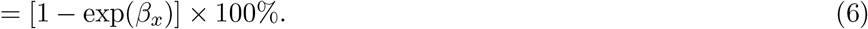

To estimate the private benefit, we calculate an analogous odds ratio relative to the odds of infection for an unvaccinated host in the same vaccinated population. For *y* ∈ {4, …18},

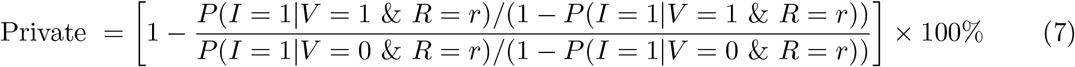

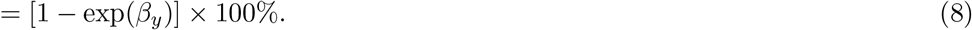

## 4 Results

### 4.1 Vaccination reduces the average amount of antigenic evolution and disease burden

For an influenza-like pathogen, vaccination reduces the average amount of antigenic evolution (Spearman’s *ρ* = −0.75, *p* < 0.001) and incidence (Spearman’s *ρ* = −0.86, *p* < 0.001, Fig. 2) when the breadth of vaccine-induced immunity is the same as that of infection. Without vaccination, the viral population evolves on average 21.5 (SD = 3.3) antigenic units and causes an average of 1.8 (SD = 0.2) cases per person over the 20-year simulation. By reducing susceptibility in the host population and the supply of beneficial mutations, vaccination decreases the number of cases and the average size of surviving mutations, thus weakening selection for antigenic novelty and increasing the strength of drift. In turn, slower antigenic evolution further reduces transmission, often driving the virus extinct. Once extinct, the viral population can no longer evolve or cause new infections. Above a 10% annual vaccination rate, implying a 28% cumulative vaccination rate over 4 years, extinction occurs rapidly, typically within 2.3 years (SD = 0.6, Fig. S7).

**Figure 2:**
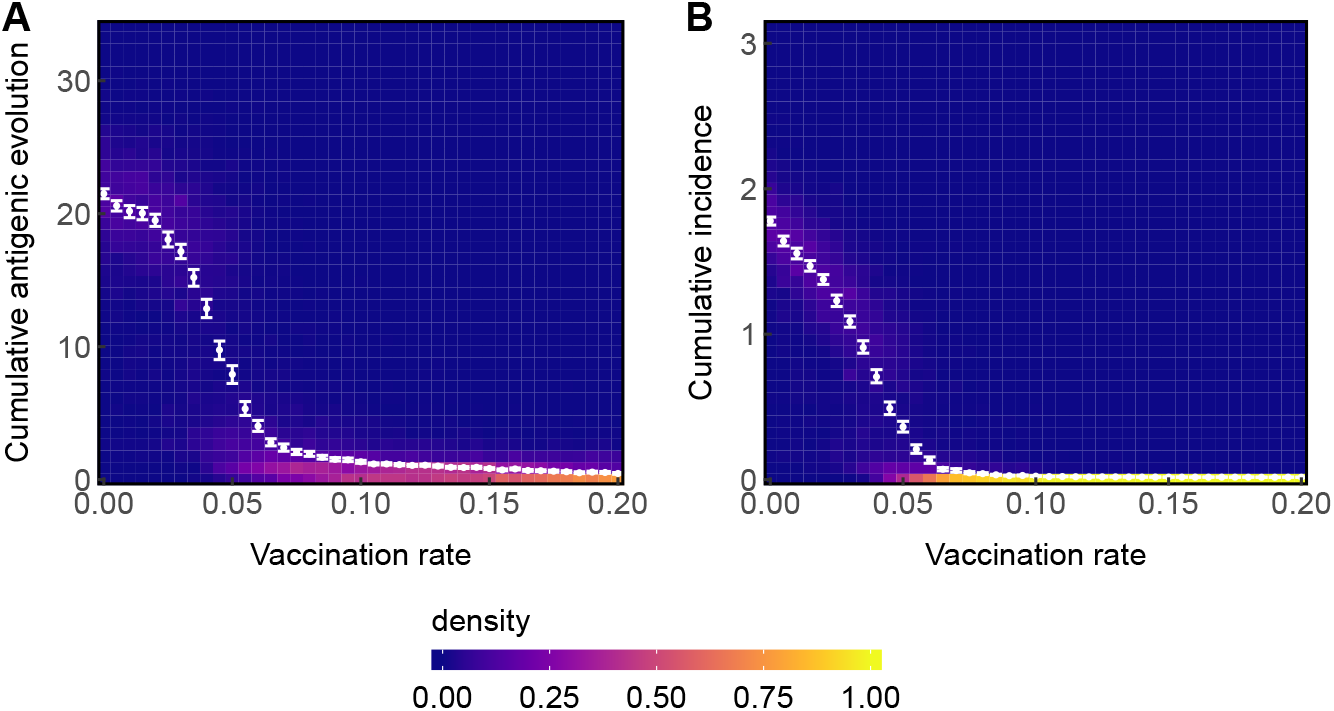
High vaccination rates decrease the average amount of (A) cumulative antigenic evolution and (B) cumulative incidence. Points show mean cumulative antigenic evolution or incidence for each vaccination rate. Error bars show 95% nonparametric bootstrapped confidence intervals of the means. Densities are calculated for each vaccination rate, such that the sum of densities for each vaccination rate equals 1. Data are collected across 500 total simulations for each rate with excessively diverse simulations (TMRCA > 10 years) excluded, leaving ~300-400 simulations per rate.

Eliminating the time interval between strain selection and vaccine distribution reduces the amount of antigenic evolution (Wilcoxon rank-sum test, *p* < 0.001) and incidence (Wilcoxon ranksum test, *p* < 0.001) even more (Fig. S8). For example, with a 300-day delay between vaccine strain selection and distribution at a 5% annual vaccination rate, the virus evolves a cumulative 7.9 (SD = 7.4) antigenic units and causes an average of 0.36 (SD =0.42) cases per person over the 20-year simulation. With zero delay at the same vaccination rate, the virus evolves a cumulative 1.4 (SD = 3.0) antigenic units and causes an average of 0.03 (SD = 0.12) cases per person over the 20-year simulation.)

Increasing the vaccination rate also decreases the probability that the viral population becomes too diverse (Fig. S9). Without vaccination, 42.5% of simulations becomes too diverse, while 35.6% and 3.4% become too diverse at a 1% and 5% annual vaccination rate, respectively. Thus, vaccination is unlikely to increase incidence by diversifying viral populations.

Given the high extinction rate with vaccination, we next examined how much these reductions in incidence could be attributed solely to the “ecological” effects of vaccination—the reduction in prevalence and increased extinction risk from accumulating herd immunity—versus the combined ecological and evolutionary impacts (Methods 3.5, Fig. S10). Relative to the case where the evolutionary effects of vaccines are blocked, vaccination with evolutionary effects can either increase or decrease both cumulative antigenic evolution and incidence (Fig. 3). Below a 3% annual vaccination rate, the virus evolves more and causes more cases when vaccination can affect evolution compared to when it cannot. The maximum difference occurs at a 1% annual vaccination rate, where the virus evolves 20.2 (95% CI 19.7-20.6) antigenic units and causes 1.6 (95% CI 1.5-1.6) cases per person over 20 years with evolutionary effects, compared to 18.7 (95% CI 18.2 – 19.2) antigenic units and 1.4 (95% CI 1.4-1.5) cases per person per 20 years without. This suggests that the virus experiences some positive selection from vaccination that buffers against slowed antigenic evolution. However, the strength of positive selection is not enough to overcome the factors that slow evolution relative to the zero vaccination case. Such factors include increased immunity against circulating strains, smaller effective viral population size reducing the probability that mutations will appear, and increased probability of stochastic extinctions due to fewer infections. The trend reverses above a 3% annual vaccination rate: at higher vaccination rates, the impact of vaccination on antigenic evolution and prevalence is much greater. Here, the maximum absolute difference occurs at a 6.5% annual vaccination rate, where the virus evolves 1.7 (95% CI 1.5-1.8) antigenic units and causes 0.033 (95% CI 0.030-0.036) cumulative cases per person with evolutionary effects, compared to 5.0 (95% CI 4.3 – 5.6) antigenic units and 0.15 (95% CI 0.13-0.18) cumulative cases per person years without.

**Figure 3:**
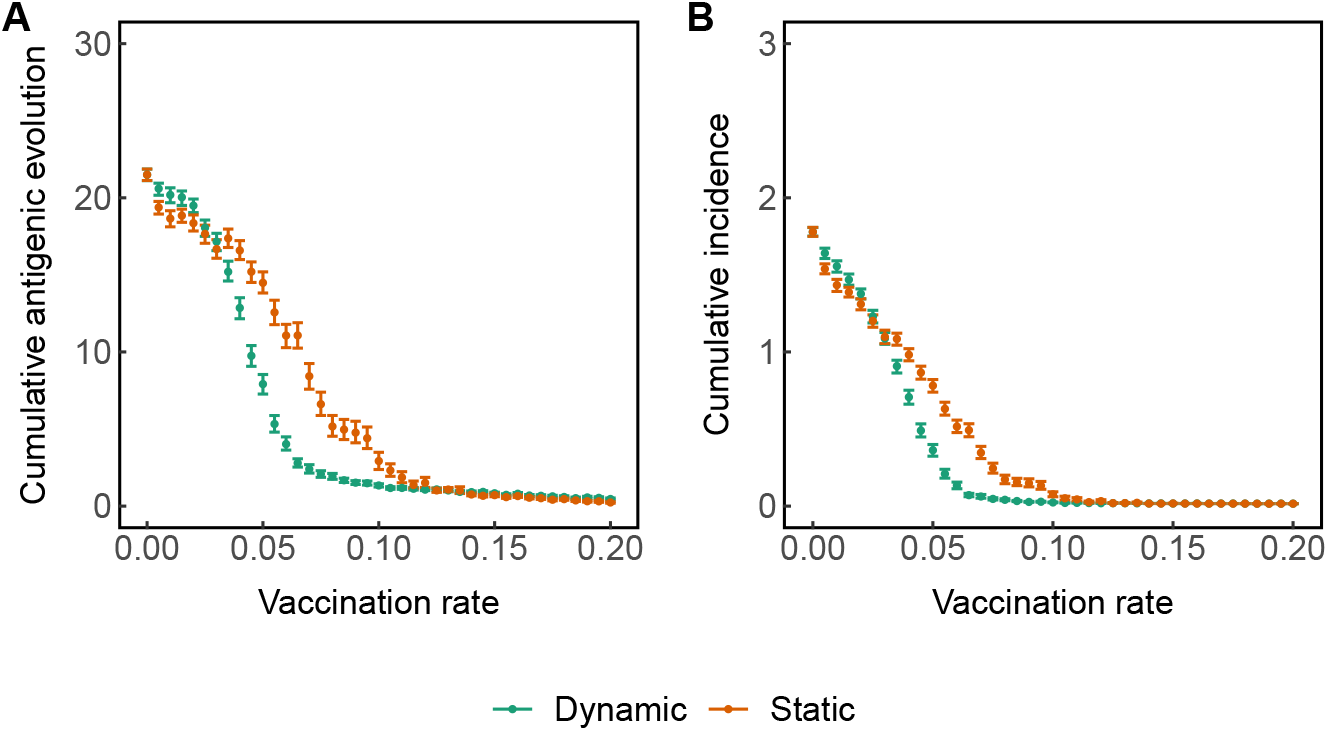
The evolutionary effects of vaccination further decrease incidence and antigenic evolution at higher vaccination rates. Green points represent simulations where vaccination can affect antigenic evolution. Orange points represent simulations where vaccination cannot affect antigenic evolution. Points show the mean cumulative (A) antigenic evolution and (B) incidence across all simulations where vaccination does (green) or does not (orange) affect antigenic evolution for each vaccination rate. Error bars show 95% nonparametric bootstrapped confidence intervals of the means. Data are collected from 500 total simulations for each vaccination rate and evolutionary condition with excessively diverse simulations excluded, leaving ~300-400 simulations per rate.

At higher vaccination rates (>3%) eradication is achieved at a lower vaccination rate when vaccination can affect antigenic evolution compared to when it cannot. For example, at an 8.5% annual vaccination rate (~20% cumulative vaccine coverage within 5 years), vaccination eradicates the virus 100% of the time (within 3.3 years on average) when vaccines can affect antigenic evolution but only does so 68% of the time (within 5.6 years on average) when vaccines cannot affect antigenic evolution (Fig. S7). This shows that when vaccination slows evolution, it does so not only by reducing the amount of evolution through extinction, but also by directly slowing evolution while the virus circulates.

The breadth of vaccine-induced immunity and the delay between vaccine strain selection and distribution change the impact of vaccination. With narrower vaccines, higher vaccination rates are needed to achieve the same average reductions in cumulative antigenic evolution and incidence using broader vaccines (Fig. S11). Regardless of breadth, distributing vaccines immediately after strain selection (i.e., distributing more antigenically matched vaccines) helps vaccines achieve the same average reductions in evolution and incidence at lower vaccination rates (Fig. S13).

### 4.2 Vaccine-driven excessive evolution is rare

We developed a statistical test to determine whether vaccination accelerates antigenic evolution or causes excessive diversity compared to the vaccine-free case. For this test, we defined excessive evolution as more than 21 antigenic units (the average amount of evolution without vaccination) over the duration of the simulation, or when the TMRCA exceeded 10 years. We counted the number of “excessively evolved” replicate simulations for each vaccination rate and breadth. If vaccination increases the rate of evolution, the frequency of excessively evolved simulations should be greater than in the vaccine-free case (Fig. S15).

We found that vaccine-driven excessive evolution only occurred at low to intermediate immune breadth (*b* = 0.2 or 0.3) and at low vaccination rates (Fig. S14). Of the parameters we tested, the most frequent cases of statistically significant vaccine-driven excessive evolution occurred at a 1.5% vaccination rate with 0.3 breadth, with 37.1% (95% CI 34.1-40.1%) of simulations showing excessive evolution (compared to 33.1% (95% CI 30.2-36.1) without vaccination). In other words, even when we detected statistically significant excessive evolution, these outcomes were at most ~ 12% more common with vaccination relative to without. However, for the influenza-like parameters considered, we conclude that vaccine-driven excessive evolution is rare.

Instances of excessive evolution are generally no more common with vaccination than without (Fig. S15). For any vaccination rate, the surviving viral populations tend to be more evolved antigenically (Fig. S11). Most of these viral populations would have evolved just as much without vaccination, and only survive vaccination *because they evolved unusually quickly.* Without vaccination, 33.1% (95% CI 30.2-36.1%) of simulations show excessive evolution. In these cases, more vaccination does not increase the rate of antigenic evolution, but instead drives slowly evolving viral populations extinct while occasionally allowing persistence of quickly evolving populations (Fig. S15). Thus, apparent increases in the amount of antigenic evolution among surviving viral populations generally reflect selection among simulations (not among viruses within a simulation) for fast-evolving populations, which appear at the same rate without vaccination.

### 4.3 Ignoring the evolutionary effects of vaccination incorrectly estimates the private and social benefits of vaccination

We next quantified the private and social benefits of vaccination to understand how ignoring evolutionary effects might bias measurements of the epidemiological effects of vaccination. We collected panel data consisting of individual hosts’ vaccination and infection histories from simulations where vaccination could affect antigenic evolution and simulations where it could not affect antigenic evolution and then fitted linear panel models to these data (Methods 3.6, Eq. 4). We define the social benefit as one minus the ratio of the odds of infection for *unvaccinated* hosts in a population vaccinated at a given rate relative to the odds in an unvaccinated population (Eq. 5). The social benefit thus measures the relative reduction in the odds of infection due to vaccination in the population. We define the private benefit as one minus the odds of infection having been vaccinated relative to the risk of infection having not been vaccinated in a population vaccinated at the given rate (Eq. 7). These metrics are the same as the direct effects (standardly reported as vaccine effectiveness [35, 55, 56]) and indirect effects of vaccination [54].

When vaccination slows antigenic evolution, we expect that the social benefit will be greater and the private benefit will be smaller than when evolutionary effects are excluded. However, although the net effect of vaccination is to slow evolution, ongoing positive selection at low vaccination rates provides buffers the decline in antigenic evolution, as described above (Fig. 3). Thus, the social (private) benefit of vaccination is not always greater (smaller) with evolutionary effects

At high vaccination rates (≥5%), the social benefit rises from vaccination’s impact on evolution. For example, when vaccination does affect antigenic evolution, at a 5% annual vaccination rate, unvaccinated hosts are 84.5% (95% CI 83.5-85.5%) less likely to be infected in a vaccinated compared to an unvaccinated population (Fig. 4, Table S3). When vaccination cannot affect antigenic evolution, an unvaccinated host in a population vaccinated at the same rate is only 61.2% (95% CI 60.2-62.1%) less likely to become infected (Fig. 4, Table S3). The same trend holds at 7% and 10% annual vaccination rates.

**Figure 4:**
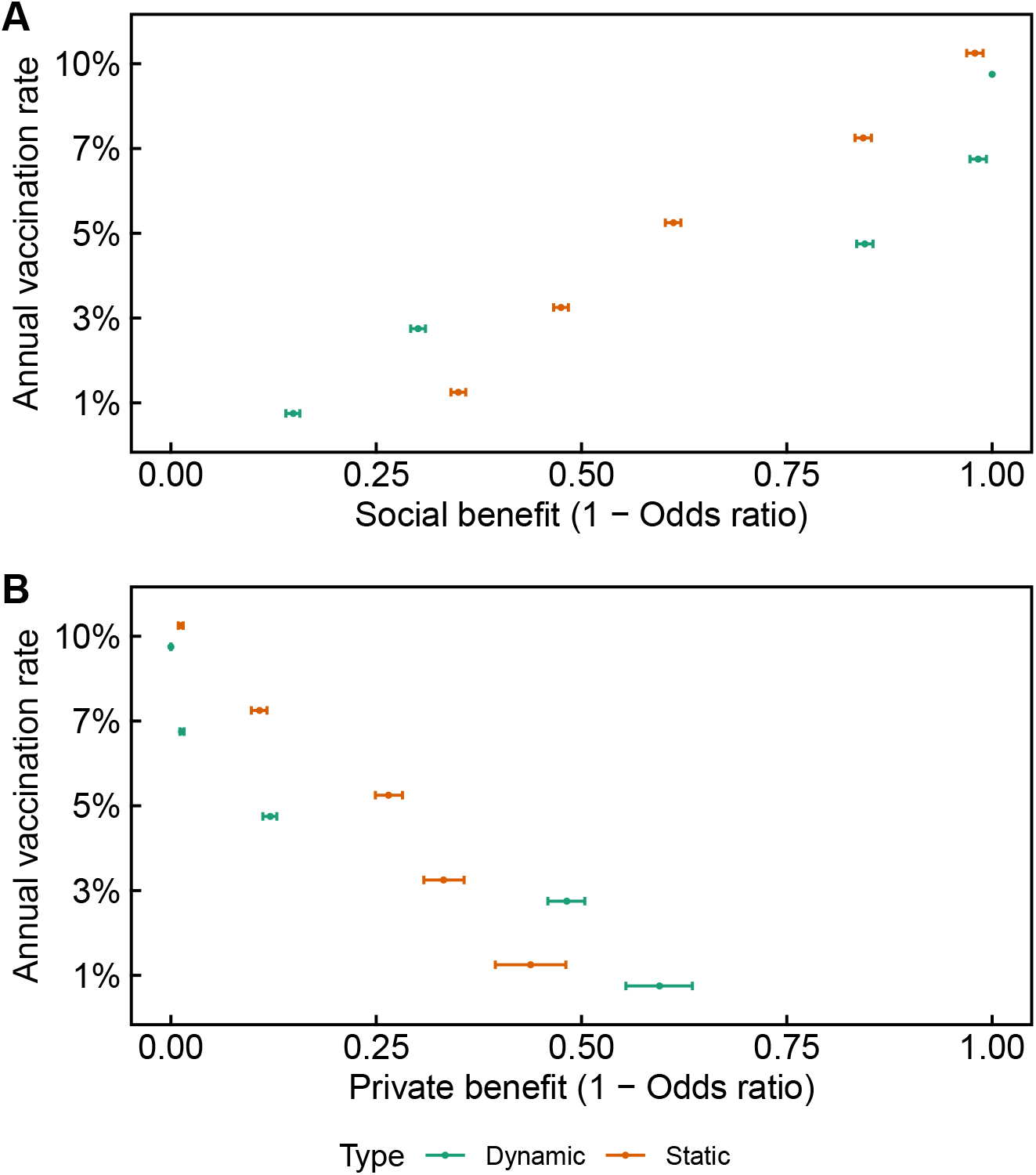
Comparison of the (A) social and (B) private benefits of vaccination when vaccination can or cannot affect antigenic evolution at 1%, 3%, 5%, 7%, and 10% annual vaccination rates. Odds ratios are calculating using coefficients from a linear panel model fitted to the last 17 years of simulated hosts’ infection and vaccination histories (Eq. 4, Table S3). Vaccination continues at the same rate after extinction, with no new infections. Mean estimates and 95% confidence intervals are shown. Green lines represent simulations where vaccination can affect antigenic evolution (dynamic). Orange lines represent simulations where vaccination cannot affect antigenic evolution (static). Points are jittered vertically for visualization.

As the social benefit rises, the private benefit falls. At a 5% annual vaccination rate, vaccinated hosts are 12.1% (95% CI 11.2-12.9%) less likely to be infected relative to unvaccinated hosts when vaccination does affect evolution, compared to 26.5% (95% CI 24.9-28.2%) less likely when vaccination does not affect evolution (Fig. 4, Table S3). Again, the same trend holds at 7% and 10% annual vaccination rates. Although in theory, improved antigenic match between the vaccine and circulating strains at high vaccination rates (Fig. S17) could increase the private benefit at higher vaccination rates, we find that the overall evolutionary impact of vaccination reduces the private benefit.

At lower vaccination rates, the trend reverses because positive selection buffers the reduction in the rate of antigenic evolution (Fig. 3): the social benefit is smaller and the private benefit is larger when vaccination can affect antigenic evolution compared to when it cannot. At a 3% annual vaccination rate, unvaccinated hosts are 30.0% (95% CI 29.3-30.7%) less likely to be infected in a vaccinated compared to an unvaccinated population when vaccination does affect antigenic evolution, compared to 47.4% (95% CI 46.9-48.0%) less likely when vaccination does not affect antigenic evolution (Fig. 4, Table S3). At the same vaccination rate, vaccinated hosts are 30.1% (95% CI 29.2-31.0%) less likely to be infected relative to unvaccinated hosts when vaccination does affect evolution, compared to 47.5% (95% CI 46.6-48.4%) less likely when vaccination does not affect evolution (Fig. 4, Table S3). The same trends hold at a 1% annual vaccination rate.

We find similar results when calculating the social benefit directly from incidence (Fig. S16). The general patterns are also similar with a vaccine that has half the breadth of natural immunity (*b* = 0.5) (Table S3).

## 5 Discussion

We found that vaccination against an H3N2-like pathogen typically slows antigenic evolution and thereby reduces disease burden beyond its immediate impact on transmission. This is a previously unrecognized potential benefit of widespread seasonal vaccination that lowers the threshold for eradication. But vaccine-induced evolution affects private and social benefits differently. At high vaccination rates, evolutionary effects increase the social benefit of vaccination and concomitantly decrease the private benefit compared to when evolutionary effects are omitted. At low vaccination rates, evolutionary effects reduce the social benefit compared to when evolutionary effects are omitted due to ongoing viral adaptation. However, the net effect of vaccination always increases the social benefit compared to without vaccination. Thus, while the evolutionary effects of vaccination may yield a large social benefit by reducing incidence as the vaccination rate increases, they may decrease the private benefit to vaccinated individuals.

The simulations’ prediction that a 10% annual vaccination rate could eradicate influenza may appear unrealistic since up to 8% of the global population is vaccinated each year [32]. To put this result in context, we highlight three key features of vaccination in the real world that differ from our model. First, vaccination is concentrated in temperate populations (e.g., the United States and Europe) rather than in the populations that contribute most to influenza’s evolution (E-S-SE Asia) [32–34]. For instance, from the 2008-2009 season to the 2014-2015 season, seasonal vaccine coverage averaged 43.4% in the United States and 13.5% across European countries, but was <1% in E-S-SE Asia [32]. Consequently, vaccination in temperate populations likely has limited impact on influenza’s persistence because the populations that sustain influenza circulation are mostly unvaccinated. Second, in contrast to our model, the same people tend to get vaccinated repeatedly, which lessens the accumulation of vaccine-induced immunity in the population over time. In the United States, up to 68.4% of vaccine recipients get vaccinated every year [57]. Third, the influenza vaccine appears to be imperfectly effective, independent of the antigenic match (as traditionally defined) between vaccine and circulating strains [56, 58, 59]. Thus, the effective amount of vaccine-induced protection in a population is probably lower than vaccine coverage estimates would suggest, especially compared to a randomly vaccinated population, implying higher vaccination rates or a more immunogenic vaccine might be necessary for eradication.

We found that the seasonal influenza vaccine is unlikely to accelerate evolution, assuming that the breadth of vaccine-induced immunity is similar to that of natural immunity. In simulations, vaccine-driven accelerated antigenic evolution only occurs when the breadth of vaccine-induced immunity is narrower than that of natural infection, and then only at low vaccination rates. The relative breadths of vaccine-induced and natural immunity are uncertain, especially since the basis of protection from infection is not precisely known. Vaccines and natural infection induce similarly broad antibody responses to the top of the hemagglutinin (i.e., as measured by serum HI), suggesting comparable breadth of immunity [60]. However, inactivated vaccines may induce fewer antibodies to neuraminidase [61], suggesting that the breadth of vaccine-induced immunity could be narrower than that of natural immunity. Host immune history also affects the generation of immune responses [62–66], and by extension the breadths of vaccine-induced and natural immunity, in ways that are largely unexplored.

Although our simulations show vaccines typically slow evolution and drive extinction in a single, closed population (i.e., a global population), other models predict faster evolution or higher incidence under different assumptions. Vaccination can accelerate antigenic evolution when stochastic extinctions in small viral populations are ignored [20]. In contrast, stochastic extinctions in our agent-based model weaken selection in small viral populations. Vaccines can also accelerate antigenic evolution locally when antigenic diversity is generated independently of vaccination, for example, when antigenic variants are introduced at a fixed rate [21,67]. In our model, strains can only emerge dynamically by mutation, so novel strains are less likely to appear when prevalence is low. In summary, the stochastic and individual-based features of our model allow for open-ended evolutionary outcomes. Mechanisms that slow down and speed up evolution interact simultaneously, with the net effect of vaccination being slower antigenic evolution.

Improved understanding of influenza’s fine-scale evolutionary and immunological dynamics might shift predictions of the impact of vaccination. For instance, the rate of vaccine-driven evolution is sensitive to transmission rates and the distribution of mutation sizes. We chose transmission and mutation parameters such that the simulated epidemiological and evolutionary dynamics match those of H3N2 [38,45]. Increasing the mutation rate, skewing the distribution of mutation sizes toward large mutations, and increasing the transmission rate each increase the rate of antigenic evolution and the tendency for viral populations to diversify [38, 45]. Our model assumes that an individual’s immune responses against multiple infections or vaccinations are independent, but immunity from prior infection or vaccination affects subsequent immune responses [68]. Consistent with this hypothesis, there is evidence that vaccination history [55,56] and recipient age (potentially a proxy for infection history) [69] affect vaccine efficacy. We also assume perfect immunogenicity, such that any reduction in vaccine efficacy is caused by antigenic mismatch. In reality, antigenic mismatch, poor immunogenicity, and poor blocking of transmission likely contribute to low efficacy, and eradication may require more vaccination than predicted by this model.

We speculate that by affecting regional antigenic evolution, vaccination has the potential to change influenza’s phylogeography. Presently, tropical and subtropical Asia contribute disproportionately to the evolution of H3N2 [33, 34], which may be due to higher regional transmission [45]. High vaccine coverage in seasonal populations may compound Asia’s propensity to produce antigenically advanced strains. Though we do not model vaccination in a metapopulation, our results suggest that vaccination in Asia might have a disproportionately large impact on influenza’s global circulation by reducing its production of antigenically advanced strains.

In theory, universal vaccines that immunize against all strains necessarily slow antigenic evolution by not discriminating between antigenic variants [19]. Our results, however, suggest that conventional seasonal influenza vaccines already have the potential to slow antigenic evolution and eradicate seasonal influenza. Increasing seasonal vaccine immunogenicity and coverage, especially in populations that contribute substantially to influenza’s evolution, could help realize similar evolutionary benefits. However, if vaccination further reduces disease burden, people may require more incentives to get vaccinated [30, 31, 70].

## 6 Data availability

The source code of the model can be found at https://github.com/cobeylab/antigen-vaccine. All data and code used to generate the results are available at https://github.com/cobeylab/vaccine-manuscript.

## 8 Author contributions

AM and SC conceived the study. FW performed the analysis and wrote the first draft of the paper. All of the authors contributed to and approved the final version.

## 9 Acknowledgements

This work was completed in part with resources provided by the University of Chicago Research Computing Center. FW and SC were supported by the National Institute of Allergy and Infectious Disease of the National Institutes of Health under award number DP2AI117921. FW was also supported by the National Institute of General Medical Sciences under award number T32GM007281. The content is solely the responsibility of the authors and does not necessarily represent the official views of the National Institutes of Health. We thank Ed Baskerville for programming guidance and Mercedes Pascual and Greg Dwyer for insightful comments.

## 1 Supplementary Information

### 1.1 Vaccination and the invasion fitness of mutants

In the following section, we develop an expectation for how vaccination affects antigenic evolution using a simple determinist model that is not related to the computational model presented in the Results. We use invasion analysis to understand how vaccination affects the invasion fitness of antigenically diverged strains by effectively reducing susceptibility. We develop an expression for the fitness of an invading mutant strain to explain how the antigenic selection gradient changes with vaccination. This preliminary analysis establishes expectations for how vaccines might affect influenza’s evolution. Unlike equation-based models, the computational agent-based model allows efficient representation of high-dimensional immune states while allowing open-ended evolution.

Here, *S*, *I*, and *R* represent the fraction of susceptible, infected, and recovered individuals. The birth rate *ν* and the death rate are equal, so the population size is constant. All individuals are born into the susceptible class. Transmission occurs at rate *β,* and recovery occurs at rate *γ*. We vaccinate some fraction *p* of newborns. In practice, this approximates vaccination of young children, who are primarily responsible for influenza transmission. Vaccinated individuals move into the recovered class.

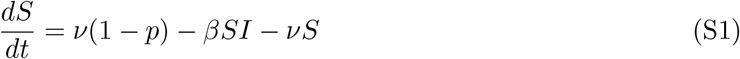

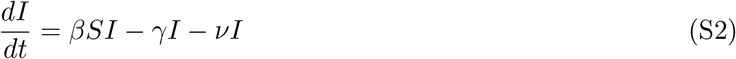

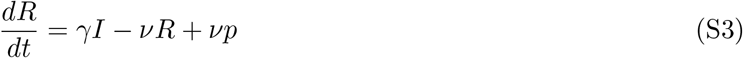

The endemic equilibrium of *S*_eq_, *I*_eq_, and *R*_eq_ is

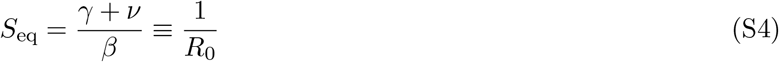

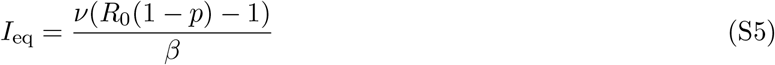

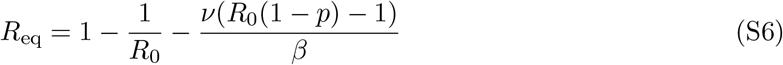

where *R*_0_, the basic reproductive number, is the number of secondary infections from a single infected individual in a totally susceptible population.

The disease-free equilibrium (when 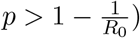) is

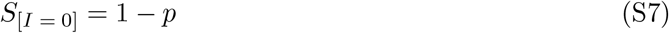

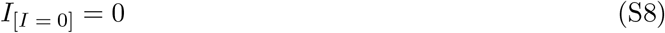

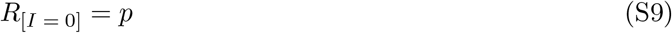

We introduce a single invading mutant 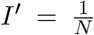, where *N* is the total population size. To find the growth rate of the mutant, we develop an expression for the amount of immunity against the mutant strain. The single mutant has an antigenic phenotype *d* antigenic units away from the resident. The conversion factor between antigenic units and infection risk is notated by *c*. Thus, the susceptibility to the mutant is given by min{*cd*, 1}, and immunity to the mutant is max{1 — *cd,* 0}. For convenience of notation, we assume *cd* ≤ 1.

We can decompose *R*_eq_ into immunity conferred by recovery from natural infection *R*_n_ and immunity conferred by vaccination *R*_v_:

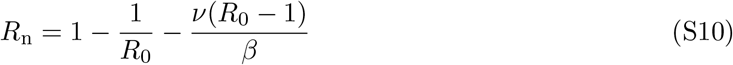

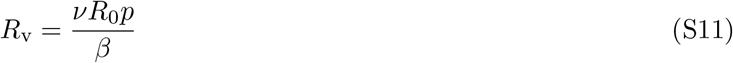

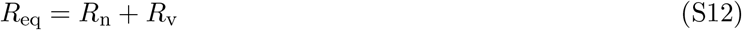

The fraction of the population immune to the invading strain is denoted *R*’. Assuming that vaccines confer a breadth of immunity relative to natural immunity *b*,

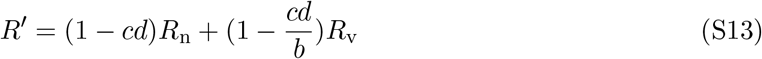

Note that when the mutant and resident are identical (*d* = 0), the immunity to the invading strain is identical to the equilibrium immunity, *R*′ = *R*_eq_. Allowing for coinfection, the fraction susceptible to the invading strain is

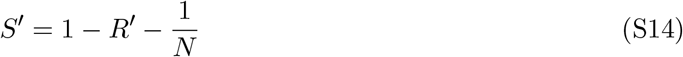

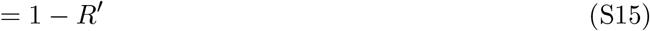

for large *N*. When the vaccination rate exceeds 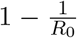, the resident is eradicated and *S*′ and *R*′ are calculated using the disease-free equilibrium.

The invasion fitness s of the mutant relative to the endemic strain is the difference between the per-capita growth rates. Note that since the resident is in equilibrium, *dI/dt* = 0.

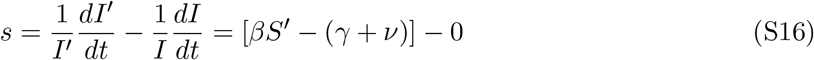

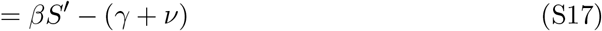

The value of *s* increases with greater distance between the mutant and resident, but decreases as more hosts become vaccinated (Fig. S1A). The expected s can be used to determine the effect of the vaccination fraction *p* on the expected invasion fitness of the mutant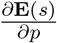. **E**(*s*) is a function of the expected distance of a mutant, **E**(*d*). In our model, we assume gamma-distributed mutation sizes with a mean *δ*_mean_ of 0.3 antigenic units and standard deviation *δ*_sd_ of 0.6 antigenic units (Fig. S1C).

**Figure S1:**
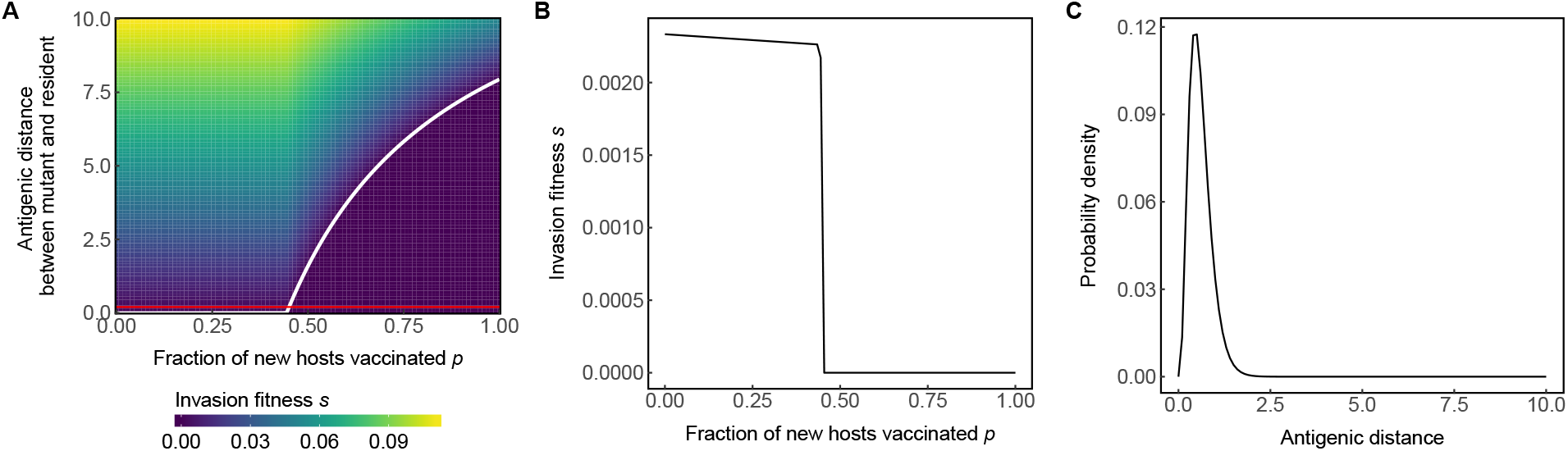
(A) High vaccination rates decrease the invasion fitness of mutant strains. For a given vaccination rate, the invasion fitness of a mutant increases with antigenic distance. However, the invasion fitness of a mutant at a given distance decreases as vaccine coverage increases. An example profile of invasion fitnesses is shown for *d* = 0.2 (the red line) in (B). Above the invasion threshold for the resident (*ρ* > 1 – 1/*R*_0_), the mutant must be increasingly more distant to invade. The white curve shows the invasion threshold, where the invasion fitness for the mutant strain is zero. Mutants above the above the curve can invade, while mutants below the curve cannot. (C) Density of gamma-distributed mutations with a *δ*_mean_ = 0.3 and *δ*_sd_ = 0.6.

We decompose 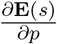 to understand how vaccines affect selection by changing susceptibility:

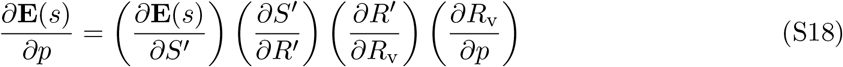

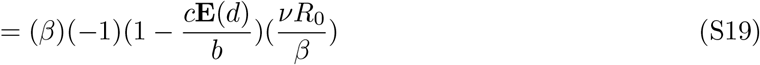

Since 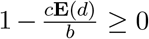 (i.e., one cannot be more than 100% immune to infection), vaccination must decrease the expected invasion fitness of the mutant, 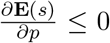, slowing evolution. This decrease is attributed to vaccination reducing susceptibility to the mutant by increasing immunity (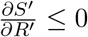 and 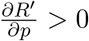) against any mutant. A larger breadth of vaccine-induced immunity (*b*) also decreases the expected invasion fitness.

### 1.2 Model validation without antigenic evolution

In the main text, we show general agreement between our simulations and observations of influenza’s epidemiology and evolution using our parameterization. We further validate the epidemiological processes of our agent-based model by removing evolution and comparing output against analytic solutions to a model using deterministic ordinary differential equations. A simple analytic solution to a model with antigenic evolution is intractable.

Classical *SIR* models include vaccination of newborns only. In a newborn-only vaccination model, the threshold eradication rate 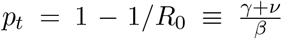. Here, we derive an eradication threshold vaccination rate for a model where all hosts are vaccinated at the same rate.

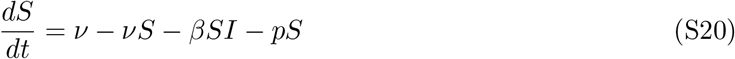

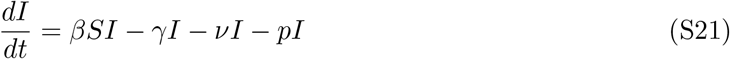

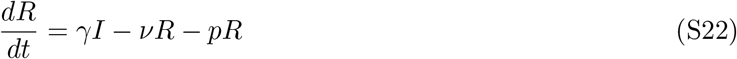

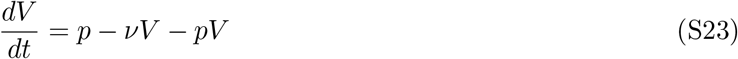

At equilibrium:

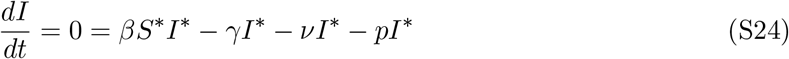

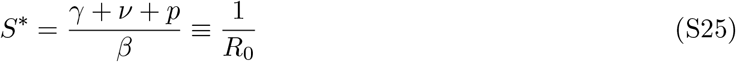

We find agreement between the simulated equilibrium fraction susceptible and the theoretical *S*\* for a range of influenza-like values of *R*_0_ (1.2-3.0) (Fig. S2).

**Figure S2:**
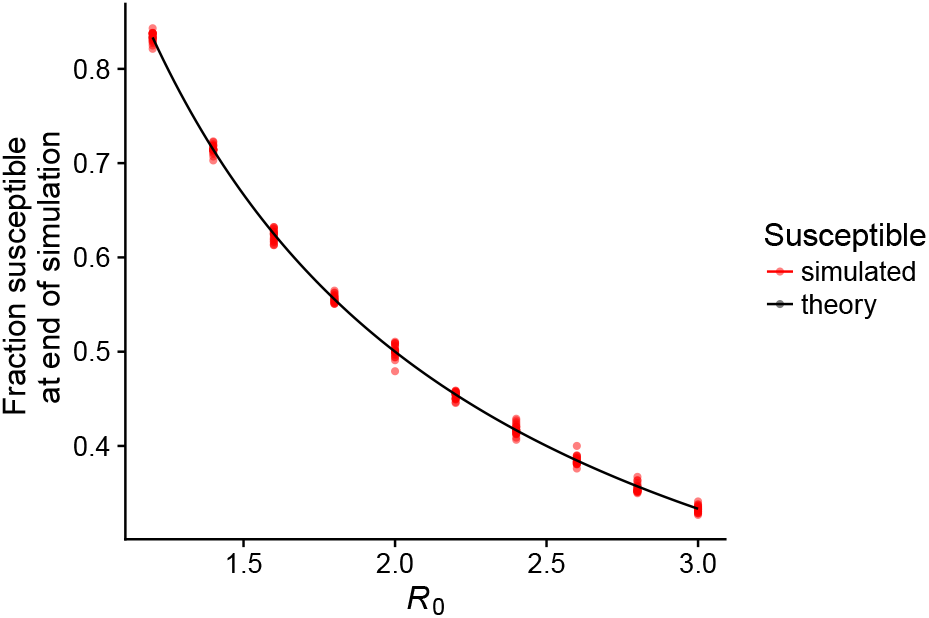
Simulated susceptible fraction at the end of 20 years without vaccination. The theoretical equilibrium fraction susceptible is given by 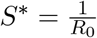. There are 40 replicate simulations shown for each value of *R*_0_.

We derive a general expression for the eradication threshold first by calculating *I*^*^:

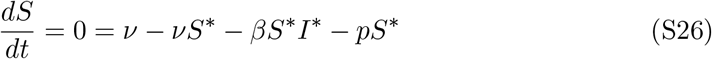

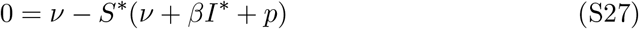

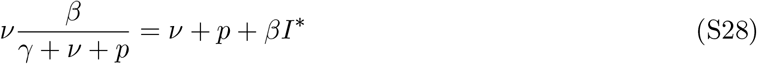

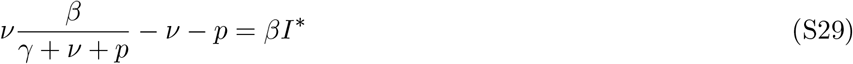

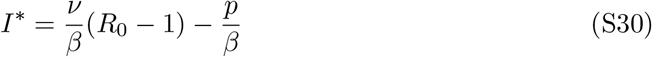

The condition for the existence of a disease-free equilibrium is *I*\* > 0. We derive an eradication threshold *p_t_* for which *I* * = 0:

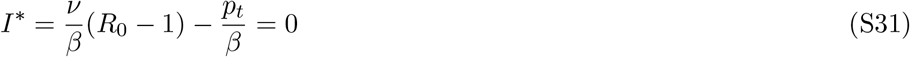

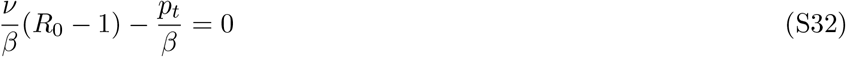

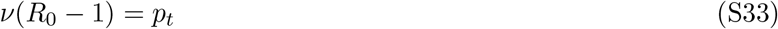

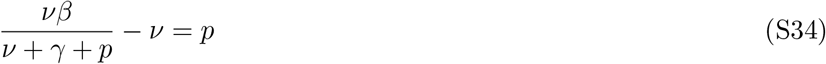

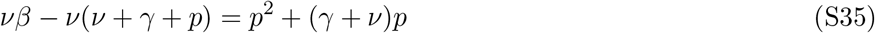

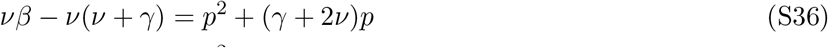

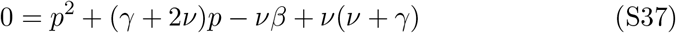

Since *p* ≥ 0, we take the nonnegative root.

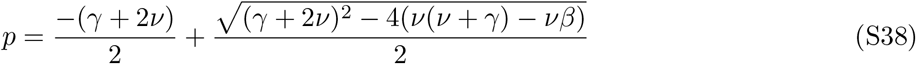

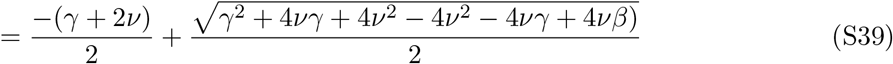

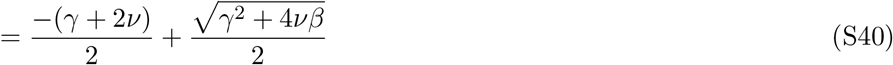

Again, we find agreement between the simulated and theoretical eradication threshold vaccination rates over a range of influenza-like values of *R*_0_ (Figs. S3, S4). Because we initialize the simulations at the endemic equilibrium *without* vaccination, some damped oscillation is to be expected, which may cause eradication at slightly lower vaccination rates than expected by theory (Fig. S5). For instance, at *R*_0_ = 1.8, theory predicts eradication at *p* = 0.0267 day^-1^, while simulation achieves extinction in 20/20 simulations within 20 years at *p* = 0.024 (Fig. S5).

The expected period of damped oscillation is derived from stability analysis. The Jacobian matrix of the SIRV model is given by

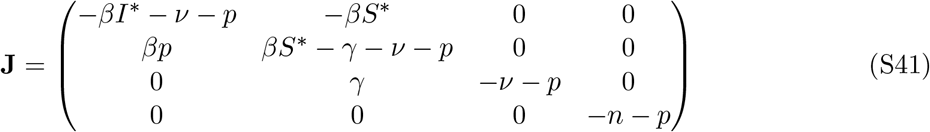

where *S*^*^ and *I*^*^ are the equilibrium fraction susceptible and infected, respectively. The period of oscillation (*T*) is inversely proportional to the imaginary part of the dominant eigenvalue of the Jacobian matrix (Λ).

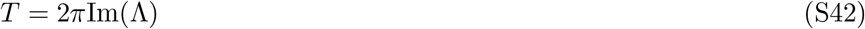

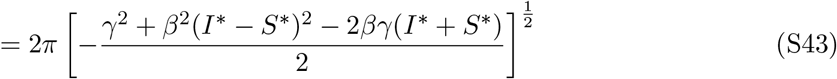

The timeseries (Fig. S5) show oscillation at annual vaccination rates of 1.3% and 1.9%. The calculated periods of oscillations at these rates are 4.5 years and 8.5 years respectively, which agree with the timeseries. Since the simulation has stochastic components, the periodicity appears more regular at first and becomes less predictable over time.

**Figure S3:**
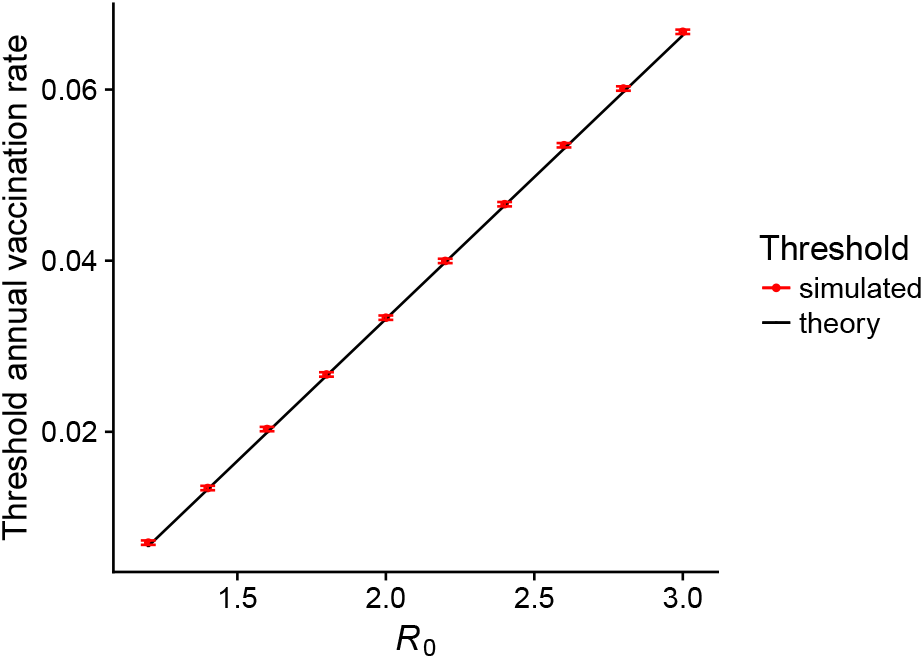
With vaccination, the simulated eradication thresholds agree with analytic predictions. The simulated threshold is the minimum vaccination rate where 40/40 simulations go extinct within 20 years. Error bars show the sampling resolution (Fig. S4). Simulations were initialized at the analytically derived equilibrium S, I, and R with vaccination (equation S40). There are 40 replicate simulations shown for each value of *R*_0_.

**Figure S4:**
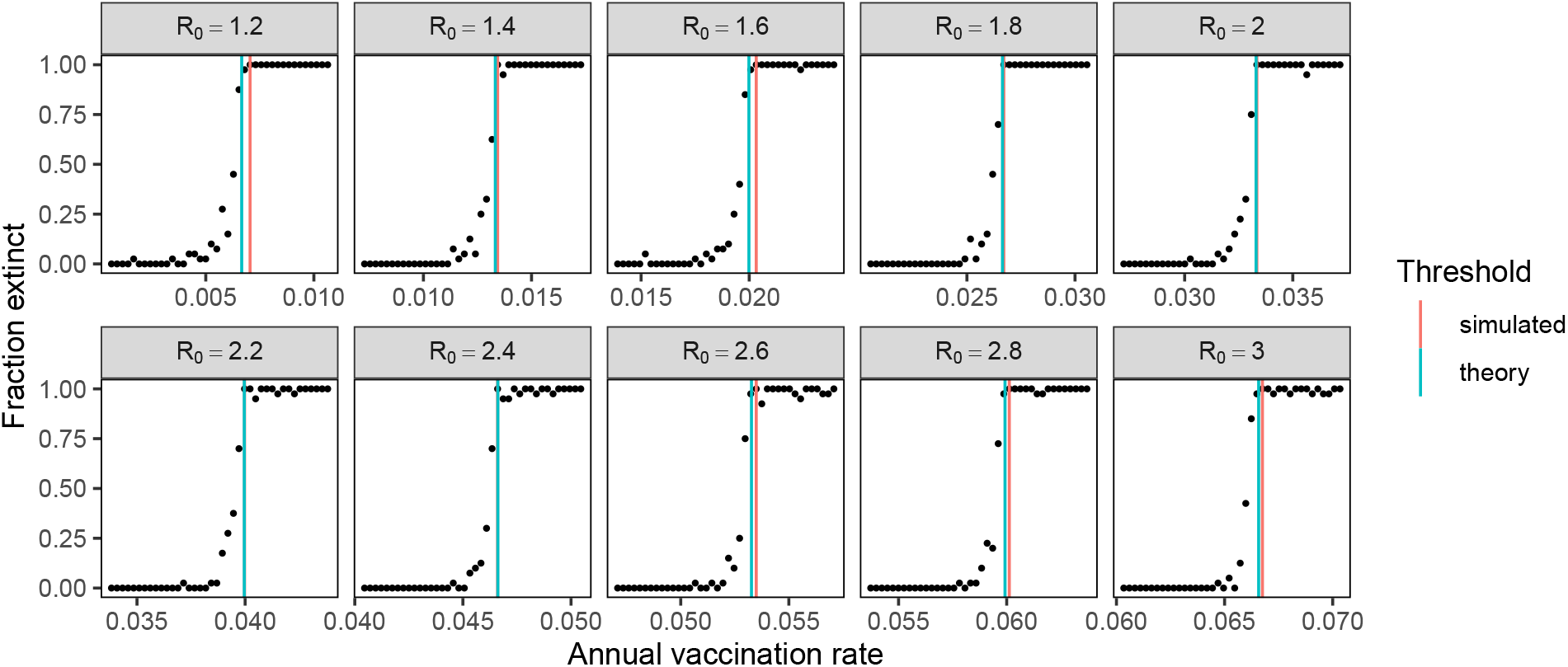
Estimation of simulated eradication thresholds without evolution, starting at the equilibrium *S, I*, and *R* with vaccination. To generate response curves, we ran 40 replicate simulations for each combination of *R*_0_ and vaccination rate and calculated the fraction of extinct simulations. The simulated eradication threshold is the minimum vaccination rate that causes 40/40 simulations to go extinct within 20 years. When the analytic equilibrium *I* was nonnegative, we initialized the simulation with a single infection.

**Figure S5:**
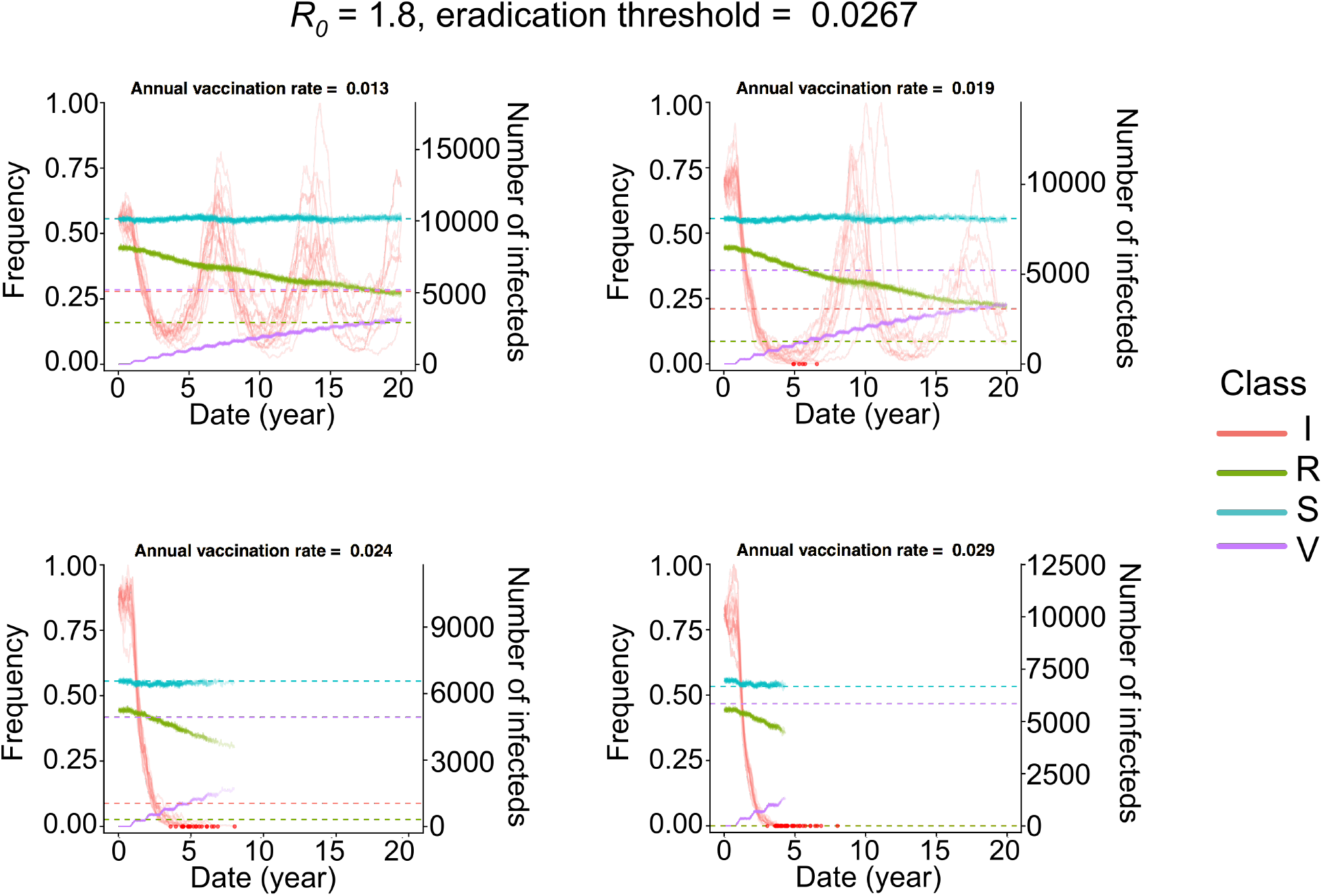
Simulated timeseries without evolution, starting at the endemic equilibrium *without* vaccination (i.e.,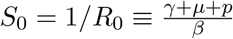, as in the manuscript, but in contrast to Appendix Figures 2 and 3). Because the population starts away from the vaccinated equilibrium, the system experiences damped oscillations, which increase the probability of stochastic extinction. Thus, we observe extinction even when the vaccination rate is slightly below the expected eradication threshold. Vaccination remains pulsed in 9-month periods, as in the model. Frequencies of susceptible (*S*), infected (*I*), recovered (*R*), and vaccinated (*V*) individuals are shown for 20 replicate simulations. The left y-axis shows the frequencies of *S* (blue), *R* (green), and *V* (purple). The right y-axis shows the number of infections (red). The dashed lines shows the expected equilibrium frequencies for each class. Red points indicate extinction events.

## 2 Supplementary tables and figures

**Table S1:** Default parameters

| Parameter | Value | Reference |
| --- | --- | --- |
| Intrinsic reproductive number ($R_0$) | 1.8 | [71, 72] |
| Duration of infection $1/\gamma$ | 5 days | [73] |
| Population size $N$ | 50 million | (see text) |
| Birth/death (turnover) rate $\nu$ | $1/30 \text{ year}^{-1}$ | [74] |
| Mutation rate $\mu$ | $10^{-4} \text{ day}^{-1}$ | (see text) |
| Mean mutation step size $\delta_{\text{mean}}$ | 0.6 antigenic units | (see text) |
| SD mutation step size $\delta_{\text{sd}}$ | 0.3 antigenic units | (see text) |
| Infection risk conversion $c$ | 0.07 | [1, 38, 44] |
| Duration of simulation | 20 years |  |
| Annual vaccination rate $r$ | 0.0-0.2 $\text{year}^{-1}$ | |
| Breadth of vaccine-induced immunity $b$ | 100% | |
| Temporal lag between vaccine strain selection and distribution $\theta$ | 300 days | |

**Table S2:** Sample panel data. Each row represents data for for individual *i* in simulation *j* at time *τ. I* is an indicator for infection status (1 if infected and 0 if not), and *V* is an indicator for vaccination status (1 if vaccinated 0 if not). *r*_1_ is an indicator for 1% vaccine coverage, *r*_5_ for 5%, and *r*_10_ for 10%.

| Identifier |  |  | Data |  |  |  |  |  |  |  | Interpretation |
| --- | --- | --- | --- | --- | --- | --- | --- | --- | --- | --- | --- |
| $\tau$ | $i$ | $j$ | $I_{ij\tau}$ | $V_{ij\tau-1}$ | $V_{ij\tau-2}$ | $V_{ij\tau-3}$ | $V_{ij\tau-4}$ | $R_{1ij}$ | $R_{5ij}$ | $R_{10ij}$ | |
| 1 | 1 | 1 | 1 | 0 | 1 | 0 | 0 | 0 | 1 | 0 | The host was infected this season (1) and only vaccinated 2 seasons ago. The population vaccination rate is 5% |
| 1 | 2 | 1 | 0 | 1 | 0 | 0 | 1 | 0 | 1 | 0 | Host not infected this season (1). Host vaccinated this season and 4 seasons ago. Population vaccination rate is 5% |
| ... |  |  |  |  |  |  |  |  |  |  |  |
| 10 | 1 | 2 | 1 | 0 | 0 | 0 | 1 | 0 | 0 | 1 | Host infected this season (10). Host vaccinated 4 seasons ago. Population vaccination rate is 10% |

**Figure S6:**
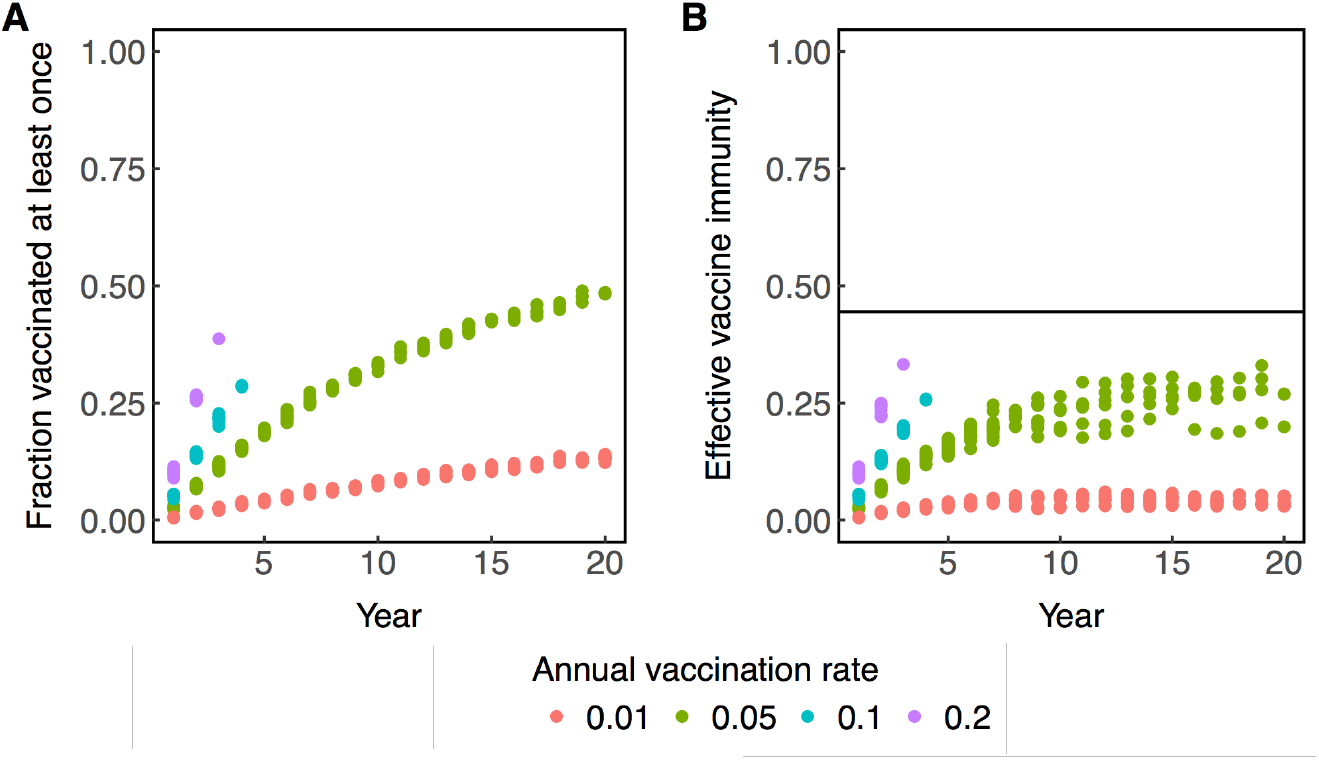
(A) Vaccine coverage and (B) effective vaccine-induced immunity over time calcu lated from simulations. (A) The fraction of individuals who have been vaccinated at least once accumulated over time and saturates at 50%. (B) The effective amount of vaccine-induced immunity in the population is calculated using the mean antigenic distance between circulating strains and the vaccinated hosts’ vaccine strains. At any given time, the effective vaccine immunity is 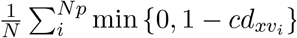, where *N* is the host population size, *p* is the fraction of vaccinated, *v_i_* is the vaccine strain received by individual *i, x* is the average circulating strain, *d* is the antigenic distance between the strains, and *c* is a constant that converts between antigenic distance and risk. The horizontal line indicates the theoretical eradication threshold in an antigenically homogenous population 1 – 1/*R*_0_.

**Figure S7:**
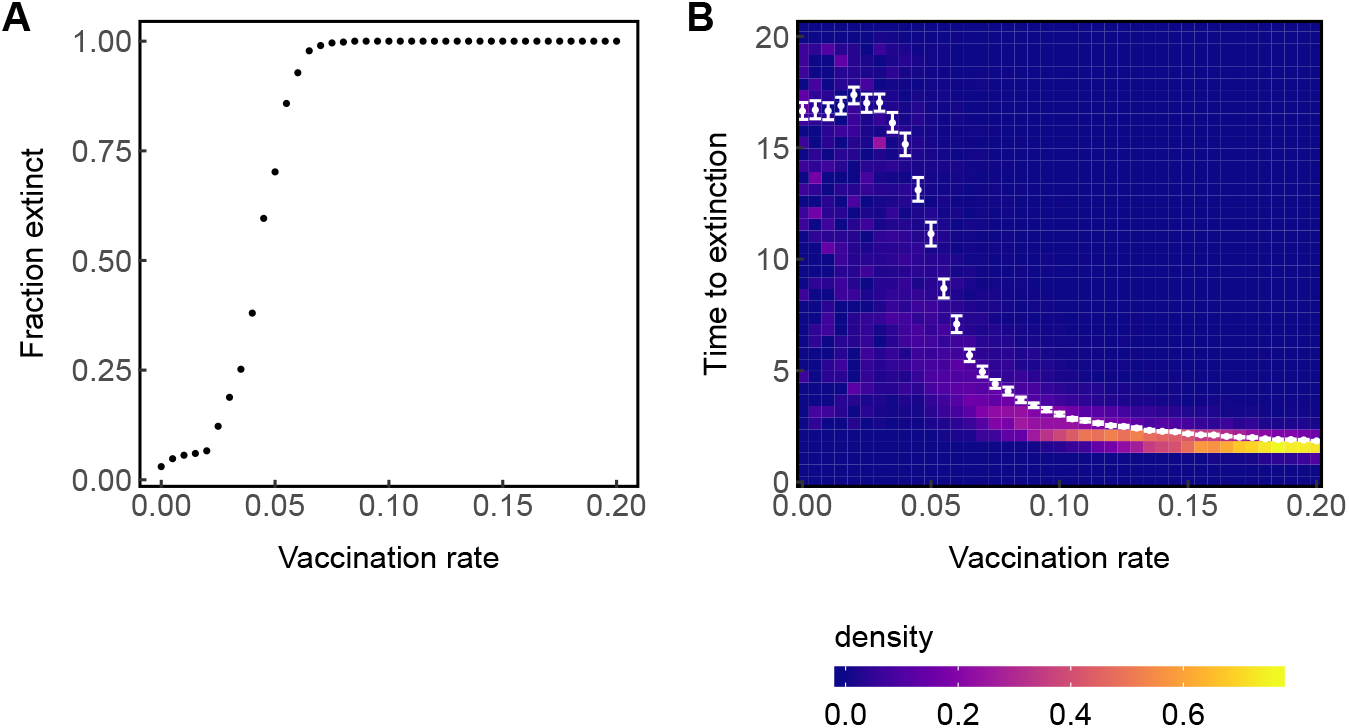
High vaccination rates increase the probability of extinction and shorten the average time to extinction. (A) Points show the fraction of simulations where the viral population went extinct before 20 years. (B) Density of times to extinction. Points show mean cumulative antigenic evolution or incidence for each vaccination rate. Error bars show 95% nonparametric bootstrapped confidence intervals of the mean. Data are collected from 500 total simulations for each vaccination rate with excessively diverse simulations (TMRCA > 10 years) excluded, leaving ~300-400 simulations per rate.

**Figure S8:**
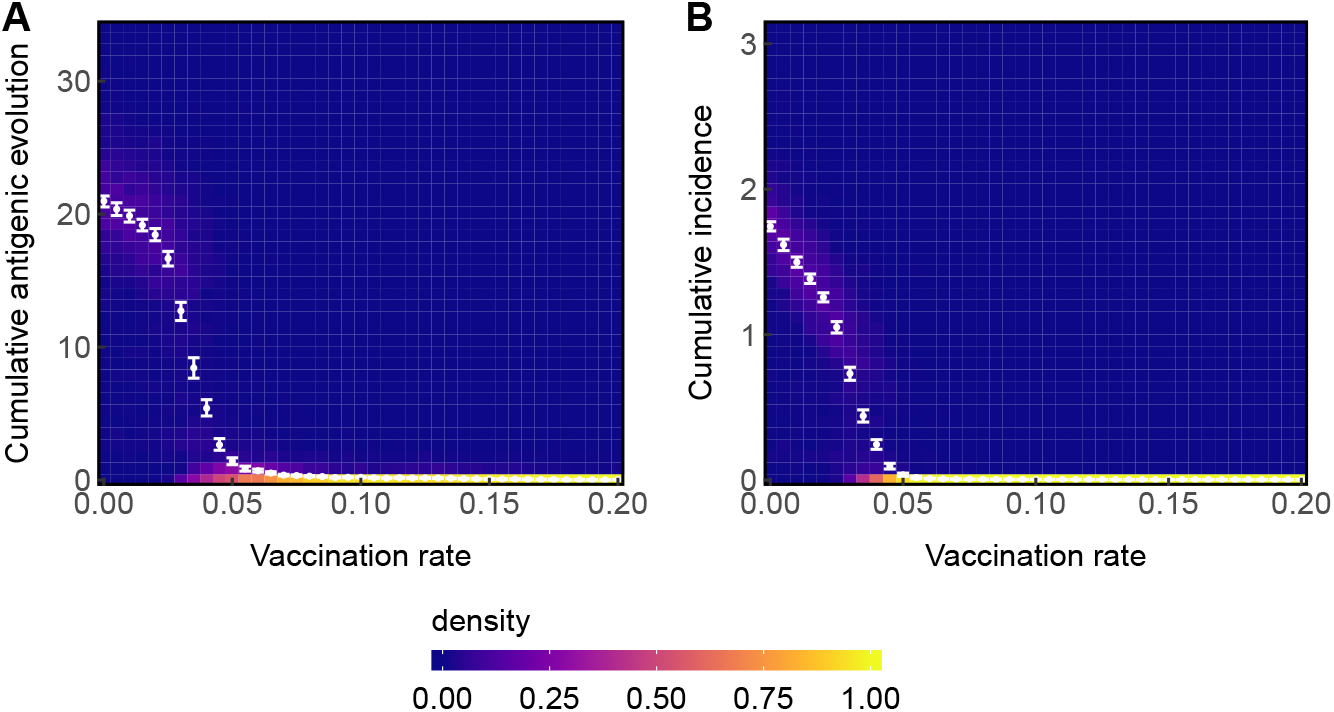
With no temporal lag between vaccine strain selection and distribution, increasing the vaccination rate quickly decreases the average amount of (A) cumulative antigenic evolution (A) and (B) incidence. Points show mean cumulative antigenic evolution or incidence for each vaccination rate. Error bars show 95% nonparametric bootstrapped confidence intervals of the mean. Data are collected from 500 total simulations for each vaccination rate with excessively diverse simulations (TMRCA > 10 years) excluded, leaving ~300-400 simulations per rate.

**Figure S9:**
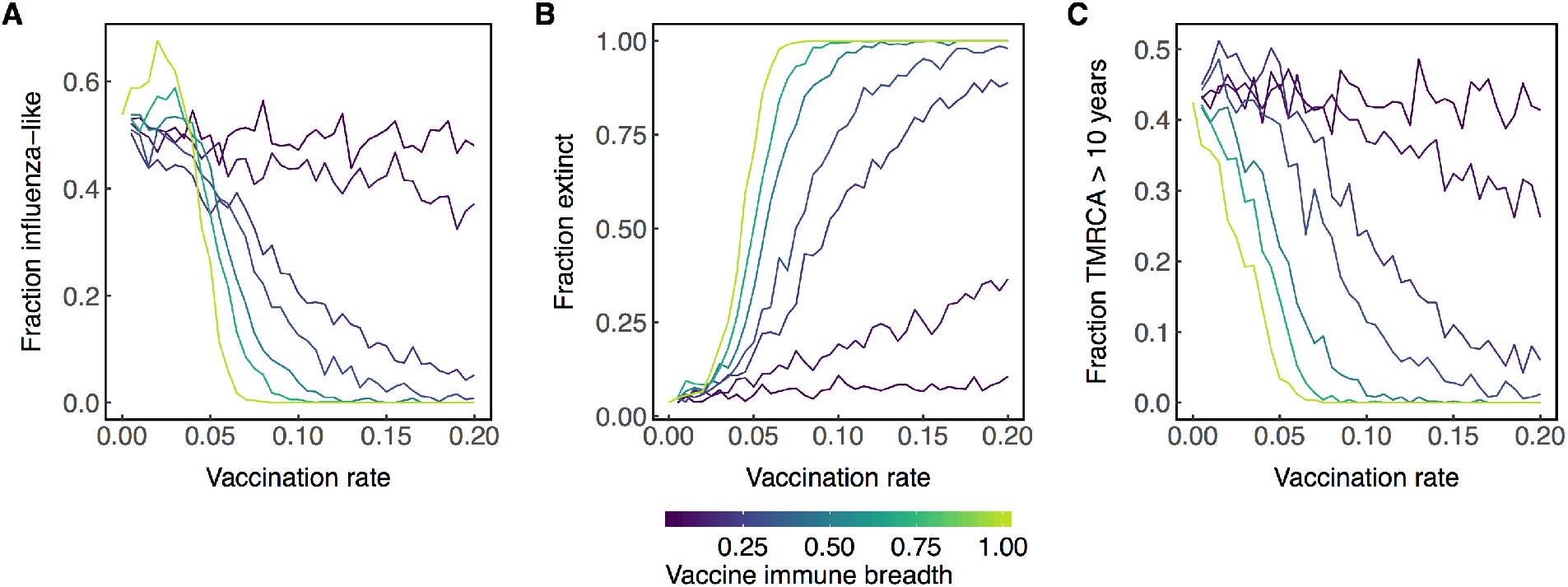
Increasing the vaccination rate increases the probability that the viral population will go extinct (B) and decreases the probability of exhibiting influenza-like dynamics (endemicity and low diversity) (A) or excessive diversification (TMRCA > 10 years) (C). Lines are colored according to the breadth of the vaccine. Data are collected from 500 replicate simulations per unique combination of vaccination rate and vaccine immune breadth with excessively diverse simulations (TMRCA > 10 years) excluded, leaving ~ 300 — 400 simulations per parameter combination.

**Figure S10:**
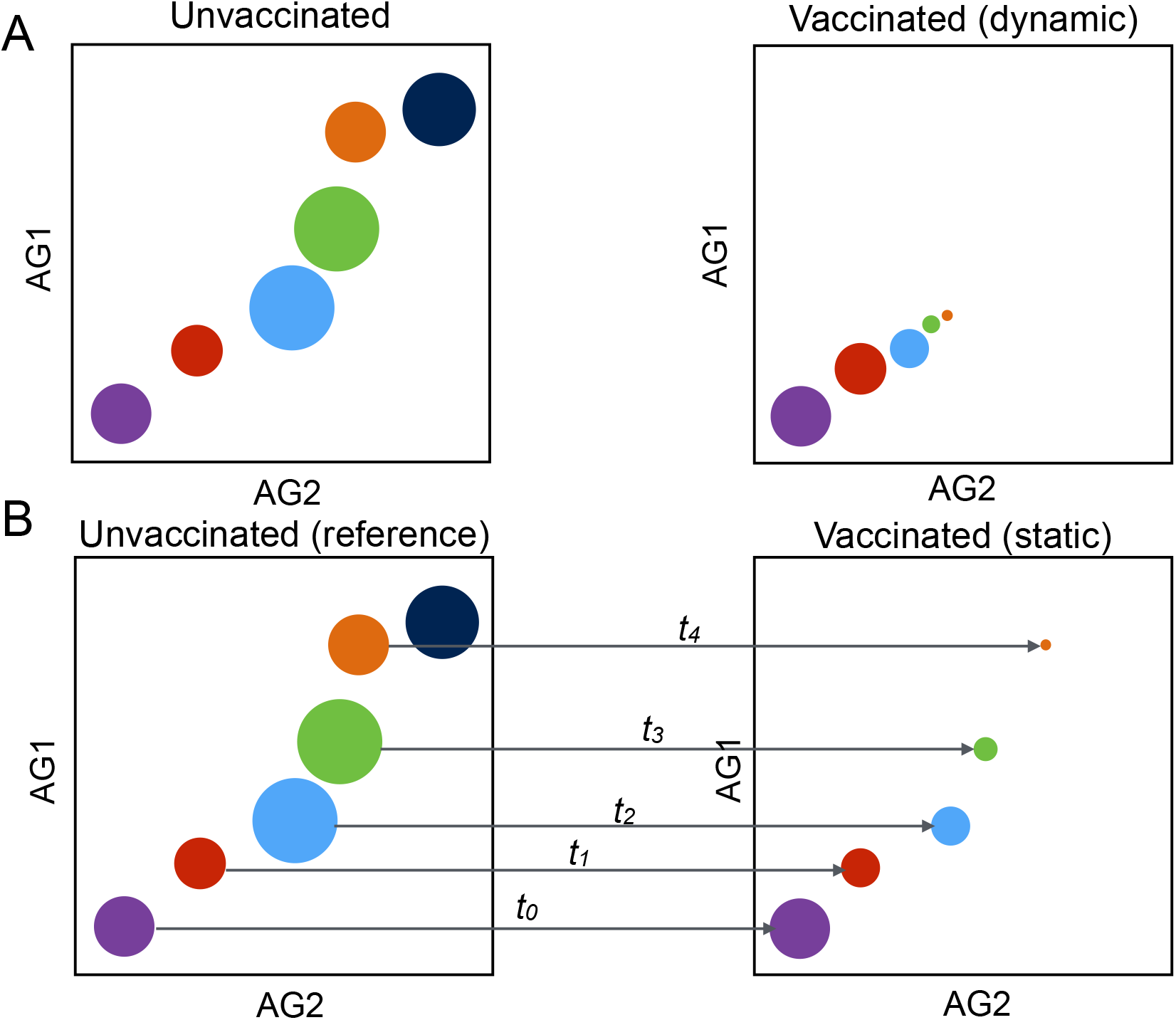
Schematic of the models where vaccination can affect antigenic evolution (dynamic, A) and where vaccination cannot affect antigenic evolution (static, B). Axes represent the principal antigenic dimensions of the 2D antigenic space. The colors of the circles represent the strain phenotypes circulating at a given time interval (the step size of the simulations is one day). The viral population starts in the lower left (purple) and evolves over time to the upper right (dark blue). The size of the circles approximates incidence. (A) In the dynamic simulations, vaccination can affect antigenic evolution. Therefore, the amount of antigenic evolution can decrease and the incidence can decrease relative to no vaccination. (B) In the static simulations, an unvaccinated population is first simulated to generate an evolutionary history that is unaffected by vaccination. Then, in the test simulation with vaccination, the antigenic phenotypes of infections during any time interval are drawn from the unvaccinated reference simulation at the contemporaneous time interval (indicated by the arrows). Thus, the rate of antigenic evolution (the position of the viral population in antigenic space at any time) is independent of vaccination. However, incidence is determined by the epidemiological dynamics of infection and recovery, so vaccination can still affect incidence (indicated by the size of the circles).

**Figure S11:**
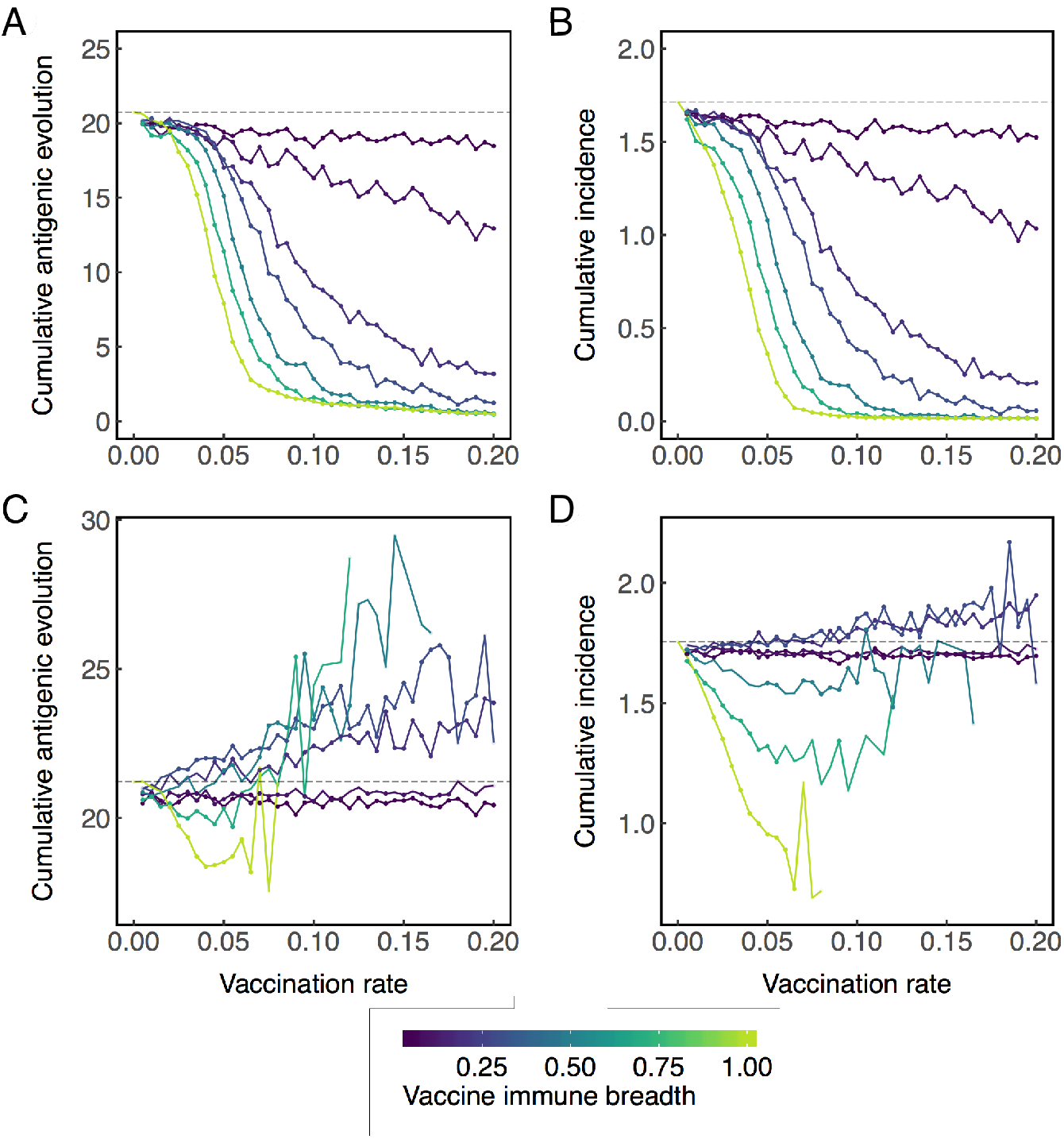
Across all simulations (A&B), vaccination decreases the average (A) cumulative antigenic evolution and (B) incidence regardless of breadth. In the subset of simulations where the viral population does not go extinct (C&D), vaccines with narrow breadth are associated with greater average antigenic evolution (C) and incidence (D), but these increases are not necessarily caused by vaccination (Fig. S14). Lines are colored according to the breadth of vaccine-induced immunity. Points indicate significant decrease (below the dashed line) or increase (above the dashed line) compared to no vaccination according to a Wilcoxon rank-sum test (*p* < 0.05) performed on at least 5 replicate simulations. Complete data are shown in Figures S12 and S15 Data are collected from 500 replicate simulations per unique combination of vaccination rate and vaccine immune breadth with excessively diverse simulations (TMRCA > 10 years) excluded, leaving ~ 300 — 400 simulations per parameter combination.

**Figure S12:**
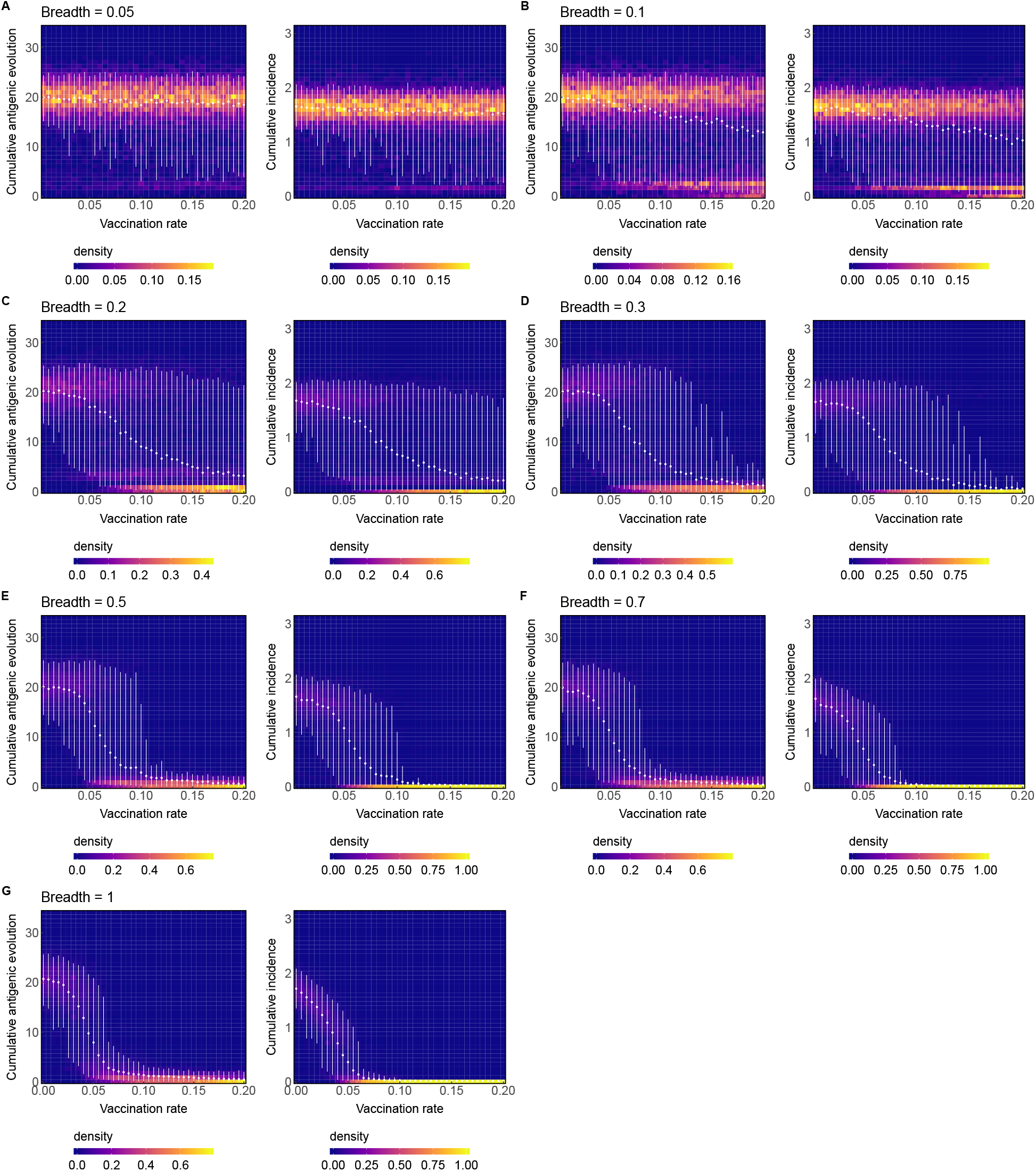
Density plots of complete simulation data corresponding to Figure S11. Points show mean cumulative antigenic evolution or incidence for each vaccination rate. Error bars show 5th and 95th percentiles for each the simulated outcomes. Data are collected from 500 replicate simulations per unique combination of vaccination rate and vaccine immune breadth with excessively diverse simulations (TMRCA > 10 years) excluded, leaving ~ 300 — 400 simulations per parameter combination.

**Figure S13:**
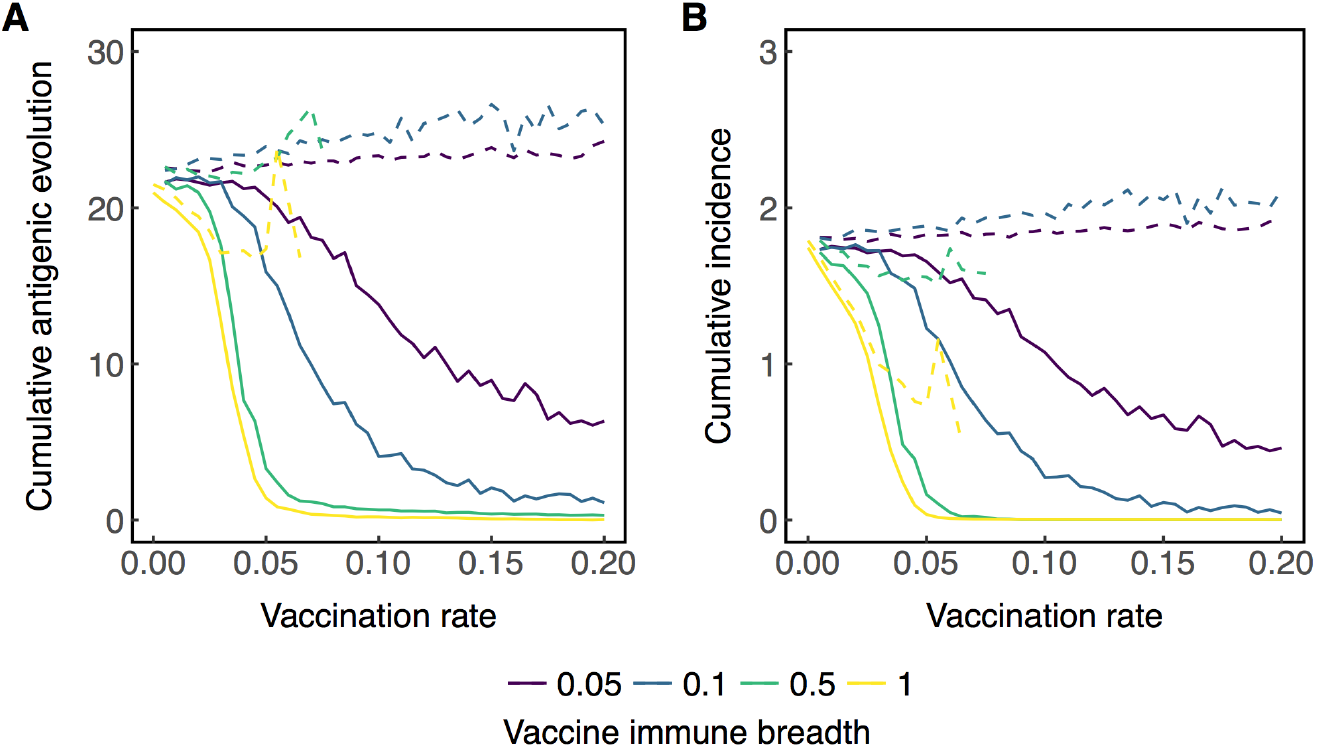
With no temporal lag between vaccine strain selection and distribution, lower vaccination rates are needed to achieve the same reductions in (A) cumulative antigenic evolution and (B) cumulative incidence compared to when vaccines are distributed 300 days after strain selection (Fig. S11). The solid lines show averages across all simulations, while dotted lines show averages over simulations where the viral population did not go extinct. Lines are colored according to the breadth of vaccine-induced immunity. Data are collected from 500 replicate simulations per unique combination of vaccination rate and vaccine immune breadth with excessively diverse simulations (TMRCA > 10 years) excluded, leaving ~ 300 — 400 simulations per parameter combination.

**Figure S14:**
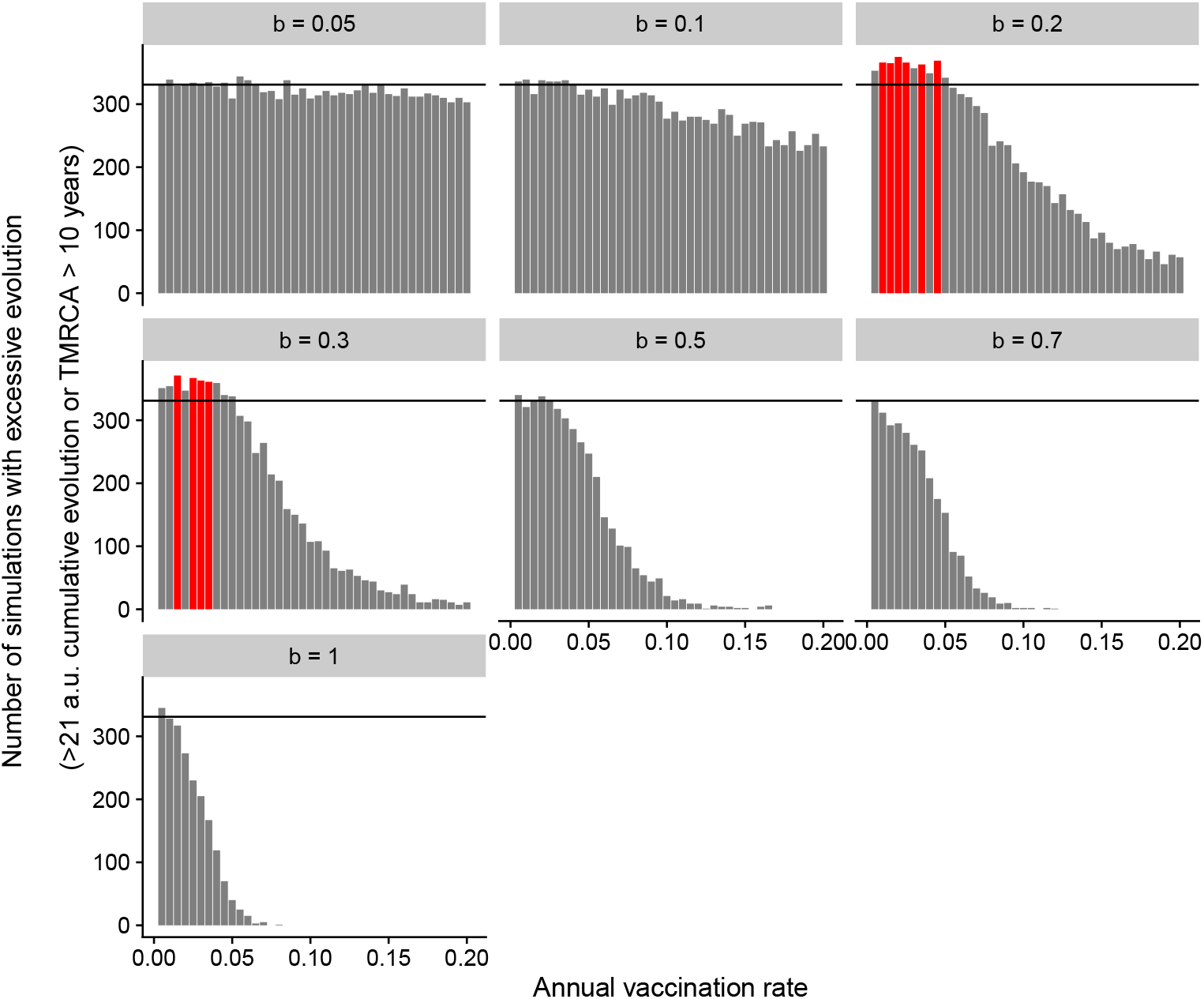
Vaccination almost always reduces the rate of antigenic evolution. The subplots show the number of simulations (out of 1000 replicates for each unique combination of parameters) that demonstrate excessive evolution for each vaccination rate and breadth *b*. Here, excessive evolution is defined by either more than 21 antigenic units of cumulative evolution or a TMRCA > 10 years. Black lines show the number of simulations that evolve excessively without vaccination (the null expectation if vaccines do not drive faster evolution). Red bars show significantly more counts of excessive evolution compared to unvaccinated simulations (*p* < 0.05, Pearson’s *χ*^2^ test).

**Figure S15:**
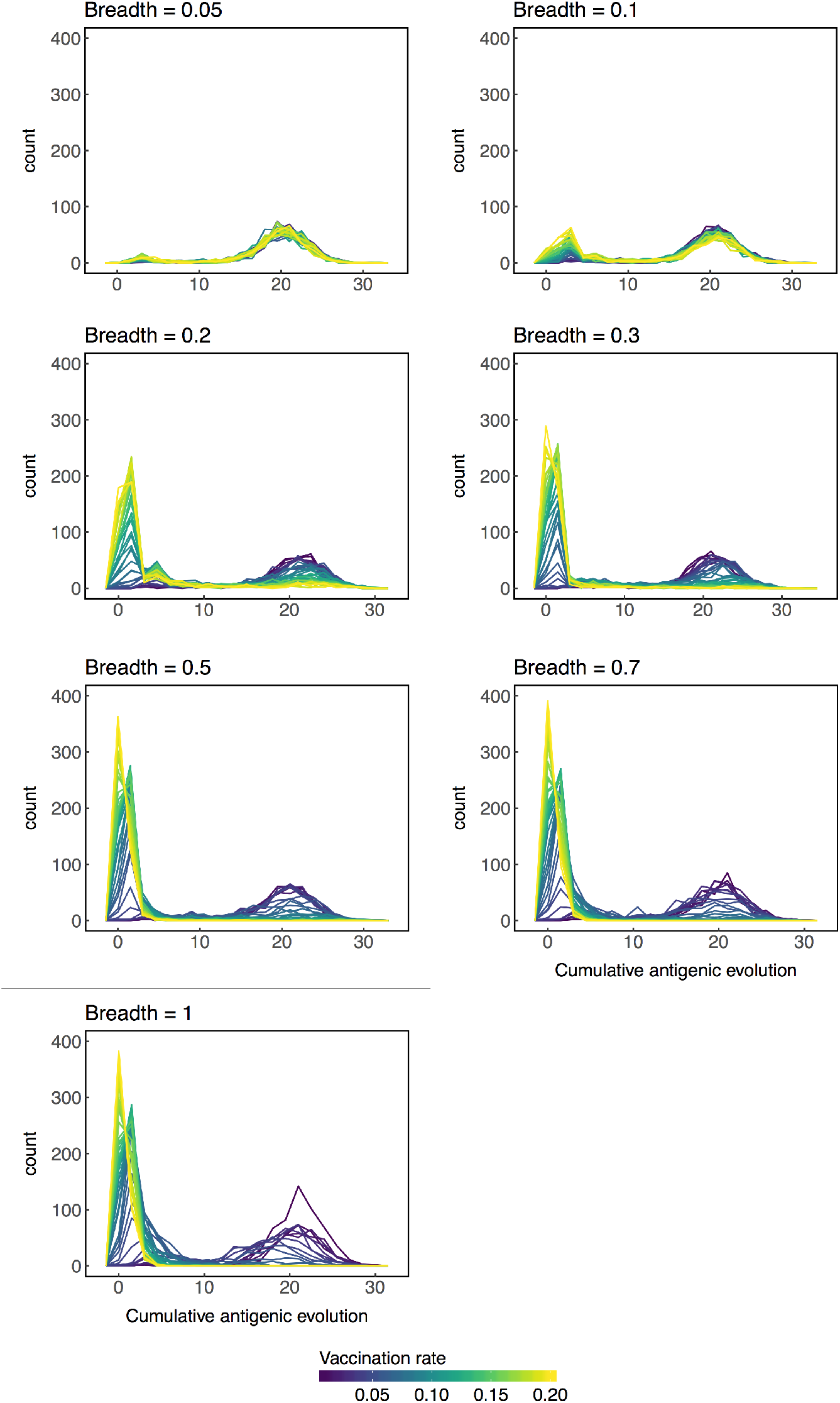
The distributions of cumulative antigenic evolution are profiles along each vaccination rate shown in figure S12. Data are collected from 500 replicate simulations per unique combination of vaccination rate and vaccine immune breadth with excessively diverse simulations (TMRCA > 10 years) excluded, leaving ~ 300 — 400 simulations per parameter combination.

**Figure S16:**
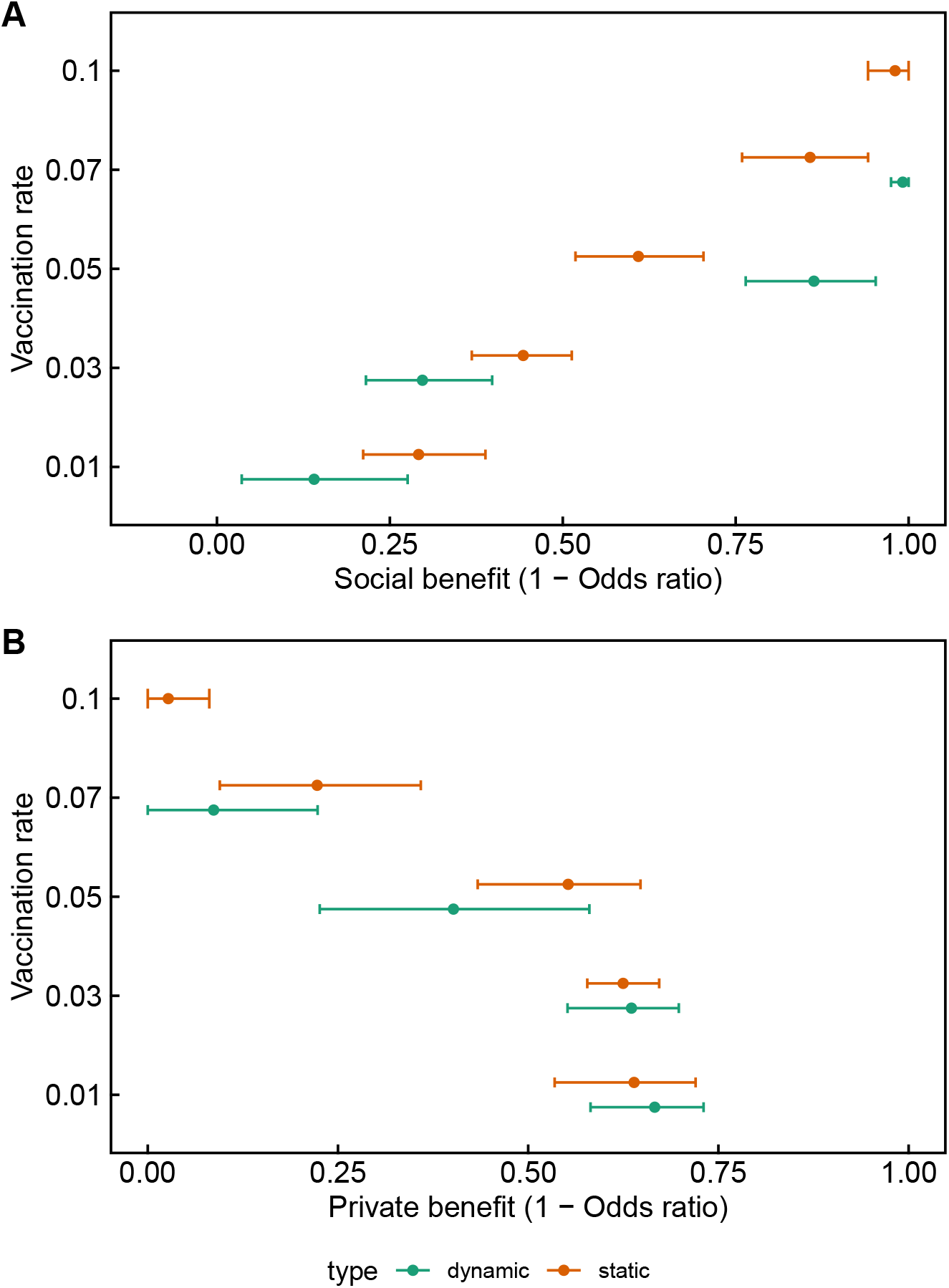
(A) Social and (B) private benefits of vaccination calculated directly from incidence as 1 — Odds ratio (Equations 5 and 7). Effects were calculated from 20 replicate simulations for each vaccination rate and simulation type using a total of 50,000 individuals for each combination of rate and simulation type. Error bars show bootstrapped 95% confidence intervals. Green lines represent simulations where vaccination can affect antigenic evolution (dynamic). Orange lines represent simulations where vaccination cannot affect antigenic evolution (static).

**Figure S17:**
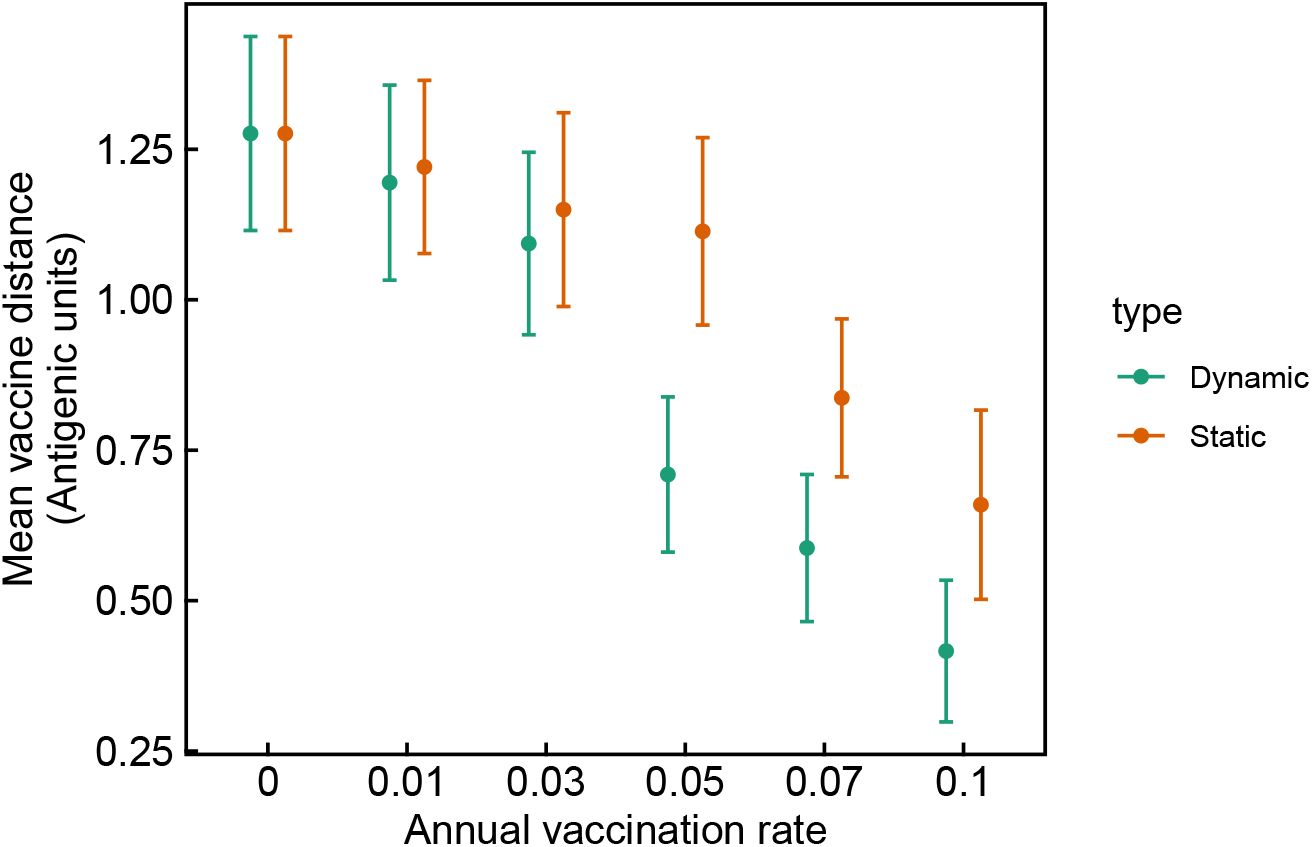
Average distance between the vaccine strain and the average antigenic phenotype of viruses circulating on the first day of the year (for the simulations used to calculate social and private benefits, Figs. 4, S16, Table S3). Distances are calculated using 20 replicate simulations for each unique vaccination rate and simulation type. Error bars show SDs. Green lines represent simulations where vaccination can affect antigenic evolution (dynamic). Orange lines represent simulations where vaccination cannot affect antigenic evolution (static).

**Table S3:** Private and social benefits of vaccination, reported as absolute risk change from linear regression. In the static model, vaccination cannot affect antigenic evolution. In the dynamic model, vaccination can affect antigenic evolution. Statistics are computed using a linear panel model on longitudinal panel data of simulated hosts’ infection and vaccination histories (equation 4). The panel data consist of 50,000 individuals per vaccination rate with 20 observations per individual. Robust standard errors shown in brackets are clustered by simulation (\**p* ≤ .05,** *p* ≤ .01,*** *p* ≤ .001)

|  | Breadth = 0.5 |  | Breadth = 1 |  |
| --- | --- | --- | --- | --- |
|  | Static | Dynamic | Static | Dynamic |
| Social benefit — 1% vac. rate | -0.026<br>(-0.027 - -0.026) | 0.009<br>(0.008 - 0.009) | -0.03<br>(-0.031 - -0.03) | -0.013<br>(-0.013 - -0.012) |
| Social benefit — 3% vac. rate | -0.036<br>(-0.037 - -0.036) | 0.01<br>(0.009 - 0.011) | -0.042<br>(-0.042 - -0.041) | -0.026<br>(-0.027 - -0.025) |
| Social benefit — 5% vac. rate | -0.053<br>(-0.054 - -0.052) | -0.031<br>(-0.031 - -0.03) | -0.054<br>(-0.055 - -0.054) | -0.077<br>(-0.077 - -0.076) |
| Social benefit — 7% vac. rate | -0.055<br>(-0.056 - -0.055) | -0.044<br>(-0.045 - -0.044) | -0.076<br>(-0.077 - -0.076) | -0.09<br>(-0.091 - -0.09) |
| Social benefit — 10% vac. rate | -0.086<br>(-0.087 - -0.085) | -0.073<br>(-0.073 - -0.072) | -0.09<br>(-0.091 - -0.09) | -0.092<br>(-0.093 - -0.092) |
| Private benefit (t-0) — 1% vac. rate | -0.021<br>(-0.026 - -0.016) | -0.039<br>(-0.044 - -0.035) | -0.038<br>(-0.042 - -0.034) | -0.053<br>(-0.056 - -0.049) |
| Private benefit (t-1) — 1% vac. rate | -0.012<br>(-0.017 - -0.006) | -0.013<br>(-0.019 - -0.007) | -0.032<br>(-0.037 - -0.028) | -0.038<br>(-0.042 - -0.034) |
| Private benefit (t-2) — 1% vac. rate | 0.004<br>(-0.003 - 0.01) | -0.01<br>(-0.016 - -0.004) | -0.021<br>(-0.026 - -0.017) | -0.028<br>(-0.033 - -0.023) |
| Private benefit (t-3) — 1% vac. rate | 0.005<br>(-0.001 - 0.011) | -0.003<br>(-0.009 - 0.003) | -0.013<br>(-0.018 - -0.007) | -0.016<br>(-0.022 - -0.011) |
| Private benefit (t-4) — 1% vac. rate | 0.005<br>(-0.001 - 0.011) | 0.002<br>(-0.005 - 0.008) | -0.009<br>(-0.014 - -0.003) | -0.005<br>(-0.011 - 0.001) |
| Private benefit (t-0) — 3% vac. rate | -0.019<br>(-0.021 - -0.016) | -0.037<br>(-0.04 - -0.034) | -0.029<br>(-0.031 - -0.027) | -0.042<br>(-0.044 - -0.04) |
| Private benefit (t-1) — 3% vac. rate | -0.009<br>(-0.012 - -0.006) | -0.021<br>(-0.024 - -0.018) | -0.023<br>(-0.025 - -0.021) | -0.034<br>(-0.036 - -0.032) |
| Private benefit (t-2) — 3% vac. rate | 0.002<br>(-0.001 - 0.006) | -0.009<br>(-0.013 - -0.006) | -0.018<br>(-0.021 - -0.016) | -0.026<br>(-0.028 - -0.023) |
| Private benefit (t-3) — 3% vac. rate | 0.003<br>(0 - 0.007) | 0<br>(-0.004 - 0.003) | -0.011<br>(-0.014 - -0.009) | -0.015<br>(-0.018 - -0.013) |
| Private benefit (t-4) — 3% vac. rate | 0.007<br>(0.004 - 0.011) | 0.002<br>(-0.002 - 0.005) | -0.006<br>(-0.009 - -0.003) | -0.012<br>(-0.015 - -0.009) |
| Private benefit (t-0) — 5% vac. rate | -0.013<br>(-0.015 - -0.011) | -0.02<br>(-0.022 - -0.018) | -0.023<br>(-0.024 - -0.021) | -0.01<br>(-0.011 - -0.01) |
| Private benefit (t-1) — 5% vac. rate | -0.005<br>(-0.007 - -0.003) | -0.009<br>(-0.011 - -0.007) | -0.016<br>(-0.018 - -0.015) | -0.008<br>(-0.009 - -0.007) |
| Private benefit (t-2) — 5% vac. rate | -0.001<br>(-0.003 - 0.001) | -0.004<br>(-0.006 - -0.001) | -0.012<br>(-0.013 - -0.01) | -0.006<br>(-0.007 - -0.005) |
| Private benefit (t-3) — 5% vac. rate | 0.002<br>(-0.001 - 0.004) | 0.001<br>(-0.001 - 0.003) | -0.009<br>(-0.01 - -0.007) | -0.004<br>(-0.005 - -0.003) |
| Private benefit (t-4) — 5% vac. rate | 0.003<br>(0.001 - 0.005) | 0.003<br>(0 - 0.005) | -0.007<br>(-0.009 - -0.005) | -0.003<br>(-0.004 - -0.002) |
| Private benefit (t-0) — 7% vac. rate | -0.016<br>(-0.017 - -0.015) | -0.014<br>(-0.015 - -0.012) | -0.009<br>(-0.01 - -0.008) | -0.001<br>(-0.001 - -0.001) |
| Private benefit (t-1) — 7% vac. rate | -0.009<br>(-0.011 - -0.008) | -0.006<br>(-0.007 - -0.004) | -0.006<br>(-0.007 - -0.005) | -0.001<br>(-0.001 - 0) |
| Private benefit (t-2) — 7% vac. rate | -0.002<br>(-0.004 - -0.001) | -0.001<br>(-0.003 - 0) | -0.005<br>(-0.006 - -0.005) | -0.001<br>(-0.001 - 0) |
| Private benefit (t-3) — 7% vac. rate | -0.001<br>(-0.002 - 0.001) | 0<br>(-0.001 - 0.002) | -0.003<br>(-0.004 - -0.002) | 0<br>(-0.001 - 0) |
| Private benefit (t-4) — 7% vac. rate | 0<br>(-0.002 - 0.002) | 0.001<br>(-0.001 - 0.002) | -0.002<br>(-0.003 - -0.001) | 0<br>(-0.001 - 0) |
| Private benefit (t-0) — 10% vac. rate | -0.002<br>(-0.002 - -0.001) | -0.003<br>(-0.004 - -0.003) | -0.001<br>(-0.001 - -0.001) | 0<br>(0 - 0) |
| Private benefit (t-1) — 10% vac. rate | -0.001<br>(-0.001 - 0) | -0.001<br>(-0.002 - 0) | -0.001<br>(-0.001 - -0.001) | 0<br>(0 - 0) |
| Private benefit (t-2) — 10% vac. rate | -0.001<br>(-0.001 - 0) | 0<br>(-0.001 - 0.001) | -0.001<br>(-0.001 - -0.001) | 0<br>(0 - 0) |
| Private benefit (t-3) — 10% vac. rate | 0<br>(-0.001 - 0.001) | 0.001<br>(0 - 0.001) | 0<br>(-0.001 - 0) | 0<br>(0 - 0) |
| Private benefit (t-4) — 10% vac. rate | 0<br>(-0.001 - 0) | 0<br>(-0.001 - 0) | 0<br>(-0.001 - 0) | 0<br>(0 - 0) |
| Constant (baseline risk) | 0.092<br>(0.091 - 0.092) | 0.092<br>(0.091 - 0.093) | 0.092<br>(0.092 - 0.093) | 0.092<br>(0.092 - 0.093) |

## References

[1] Gupta V, Earl DJ, Deem MW. 2006 Quantifying influenza vaccine efficacy and antigenic distance. Vaccine 24, 3881–3888.

[2] Bridges CB, Thompson WW, Meltzer MI, Reeve GR, Talamonti WJ, Cox NJ, Lilac HA, Hall H, Klimov A, Fukuda K. 2000 Effectiveness and cost-benefit of influenza vaccination of healthy working adults: A randomized controlled trial. JAMA 284, 1655–1663.

[3] Belongia EA, Kieke BA, Donahue JG, Greenlee RT, Balish A, Foust A, Lindstrom S, Shay DK, Marshfield Influenza Study Group. 2009 Effectiveness of inactivated influenza vaccines varied substantially with antigenic match from the 2004-2005 season to the 2006-2007 season. The Journal of Infectious Diseases 199, 159–167.

[4] Zimmerman RK, Nowalk MP, Chung J, Jackson ML, Jackson LA, Petrie JG, Monto AS, McLean HQ, Belongia EA, Gaglani M, Murthy K, Fry AM, Flannery B. 2016 2014-2015 Influenza Vaccine Effectiveness in the United States by Vaccine Type. Clinical Infectious Diseases 63, 1564–1573.

[5] Carrat F, Flahault A. 2007 Influenza vaccine: The challenge of antigenic drift. Vaccine 25, 6852–6862.

[6] Kennedy DA, Read AF. 2017 Why does drug resistance readily evolve but vaccine resistance does not?. Proceedings. Biological sciences 284, 20162562.

[7] Kennedy DA, Read AF. 2018 Why the evolution of vaccine resistance is less of a concern than the evolution of drug resistance. Proceedings of the National Academy of Sciences 115, 12878–12886.

[8] Lee CW, Senne DA, Suarez DL. 2004 Effect of Vaccine Use in the Evolution of Mexican Lineage H5N2 Avian Influenza Virus. Journal of Virology 78, 8372–8381.

[9] Read AF, Baigent SJ, Powers C, Kgosana LB, Blackwell L, Smith LP, Kennedy DA, Walkden-Brown SW, Nair VK. 2015 Imperfect Vaccination Can Enhance the Transmission of Highly Virulent Pathogens. PLOS Biology 13, e1002198.

[10] Weinberger DM, Malley R, Lipsitch M. 2011 Serotype replacement in disease after pneumococcal vaccination. The Lancet 378, 1962–1973.

[11] Wen F, Bell S, Bedford T, Cobey S, Wen FT, Bell SM, Bedford T, Cobey S. 2018 Estimating Vaccine-Driven Selection in Seasonal Influenza. Viruses 10, 509.

[12] Bansal S, Pourbohloul B, Meyers LA. 2006 A comparative analysis of influenza vaccination programs. PLoS Medicine 3, e387–10.

[13] Weycker D, Edelsberg J, Halloran ME, Longini IM, Nizam A, Ciuryla V, Oster G. 2005 Population-wide benefits of routine vaccination of children against influenza. Vaccine 23, 1284–1293.

[14] Medlock J, Galvani AP. 2009 Optimizing influenza vaccine distribution. Science 325, 1705–1708.

[15] Shim E, Galvani AP. 2012 Distinguishing vaccine efficacy and effectiveness. Vaccine 30, 6700–6705.

[16] Arinaminpathy N, Kim IK, Gargiullo P, Haber M, Foppa IM, Gambhir M, Bresee J. 2017 Estimating Direct and Indirect Protective Effect of Influenza Vaccination in the United States. American journal of epidemiology 186, 92–100.

[17] Gog JR, Grenfell BT. 2002 Dynamics and selection of many-strain pathogens. Proceedings of the National Academy of Sciences 99, 17209–17214.

[18] Kucharski A, Gog JR. 2012 Influenza emergence in the face of evolutionary constraints. Proceedings of the Royal Society B 279, 645–652.

[19] Arinaminpathy N, Ratmann O, Koelle K, Epstein SL, Price GE, Viboud C, Miller MA, Grenfell BT. 2012 Impact of cross-protective vaccines on epidemiological and evolutionary dynamics of influenza. Proceedings of the National Academy of Sciences of the United States of America 109, 3173–3177.

[20] Boni MF, Gog JR, Andreasen V, Feldman MW. 2006 Epidemic dynamics and antigenic evolution in a single season of influenza A. Proceedings of the Royal Society B 273, 1307–1316.

[21] Subramanian R, Graham AL, Grenfell BT, Arinaminpathy N. 2016 Universal or Specific? A Modeling-Based Comparison of Broad-Spectrum Influenza Vaccines against Conventional, Strain-Matched Vaccines. PLoS Computational Biology 12, e1005204–17.

[22] Kennedy DA, Dunn PA, Read AF. 2018 Modeling Marek’s disease virus transmission: A framework for evaluating the impact of farming practices and evolution. Epidemics 23, 85–95.

[23] Demicheli V, Jefferson T, Ferroni E, Rivetti A, Di Pietrantonj C. 2018 Vaccines for preventing influenza in healthy adults. Cochrane Database of Systematic Reviews 2, CD001269.

[24] Hurwitz ES, Haber M, Chang A, Shope T, Teo S, Ginsberg M, Waecker N, Cox NJ. 2000 Effectiveness of influenza vaccination of day care children in reducing influenza-related morbidity among household contacts. JAMA 284, 1677–1682.

[25] Principi N, Esposito S, Marchisio P, Gasparini R, Crovari P. 2003 Socioeconomic impact of influenza on healthy children and their families. The Pediatric Infectious Disease Journal 22, S207–10.

[26] Loeb M, Russell ML, Moss L, Fonseca K, Fox J, Earn DJD, Aoki F, Horsman G, Van Caeseele P, Chokani K, Vooght M, Babiuk L, Webby R, Walter SD. 2010 Effect of influenza vaccination of children on infection rates in Hutterite communities: a randomized trial. JAMA 303, 943–950.

[27] Pebody RG, Green HK, Andrews N. 2015 Uptake and impact of vaccinating school age children against influenza during a season with circulation of drifted influenza A and B strains, England, 2014/15. Euro surveillance 20, 1560–7917.

[28] Charu V, Viboud C, Simonsen L, Sturm-Ramirez K, Shinjoh M, Chowell G, Miller M, Sugaya N. 2011 Influenza-Related Mortality Trends in Japanese and American Seniors: Evidence for the Indirect Mortality Benefits of Vaccinating Schoolchildren. PLoS ONE 6, e26282.

[29] Geoffard PY, Philipson T. 1997 Disease Eradication: Private versus Public Vaccination. The American Economic Review 87, 222–230.

[30] Brewer NT, Chapman GB, Gibbons FX, Gerrard M, McCaul KD, Weinstein ND. 2007 Metaanalysis of the relationship between risk perception and health behavior: The example of vaccination. Health Psychology 26, 136–145.

[31] Chapman GB, Coups EJ. 1999 Predictors of influenza vaccine acceptance among healthy adults. Preventive medicine 29, 249–262.

[32] Palache A, Oriol-Mathieu V, Fino M, Xydia-Charmanta M, Influenza Vaccine Supply task force (IFPMA IVS). 2015 Seasonal influenza vaccine dose distribution in 195 countries (2004-2013): Little progress in estimated global vaccination coverage. Vaccine 33, 5598–5605.

[33] Bedford T, Cobey S, Beerli P, Pascual M. 2010 Global migration dynamics underlie evolution and persistence of human influenza A (H3N2). PLoS Pathogens 6, e1000918.

[34] Bedford T, Riley S, Barr IG, Broor S, Chadha M, Cox NJ, Daniels RS, Gunasekaran CP, Hurt AC, Kelso A, Klimov A, Lewis NS, Li X, McCauley JW, Odagiri T, Potdar V, Rambaut A, Shu Y, Skepner E, Smith DJ, Suchard MA, Tashiro M, Wang D, Xu X, Lemey P, Russell CA. 2015 Global circulation patterns of seasonal influenza viruses vary with antigenic drift. Nature 523, 217–220.

[35] Osterholm MT, Kelley NS, Sommer A, Belongia EA. 2012 Efficacy and effectiveness of influenza vaccines: a systematic review and meta-analysis. The Lancet. Infectious diseases 12, 36–44.

[36] Belongia EA, Simpson MD, King JP, Sundaram ME, Kelley NS, Osterholm MT, McLean HQ. 2016 Variable influenza vaccine effectiveness by subtype: a systematic review and meta-analysis of test-negative design studies. The Lancet. Infectious diseases 16, 942–951.

[37] Tam JS, Capeding MRZ, Lum LCS, Chotpitayasunondh T, Jiang Z, Huang LM, Lee BW, Qian Y, Samakoses R, Lolekha S, Rajamohanan KP, Narayanan SN, Kirubakaran C, Rappaport R, Razmpour A, Gruber WC, Forrest BD, Pan-Asian CAIV-T Pediatric Efficacy Trial Network. 2007 Efficacy and safety of a live attenuated, cold-adapted influenza vaccine, trivalent against culture-confirmed influenza in young children in Asia. The Pediatric Infectious Disease Journal 26, 619–628.

[38] Bedford T, Rambaut A, Pascual M. 2012 Canalization of the evolutionary trajectory of the human influenza virus. BMC Biology 10, 38.

[39] Smith DJ, Lapedes AS, de Jong JC, Bestebroer TM, Rimmelzwaan GF, Osterhaus ADME, Fouchier RAM. 2004 Mapping the antigenic and genetic evolution of influenza virus. Science 305, 371–376.

[40] Bedford T, Suchard MA, Lemey P, Dudas G, Gregory V, Hay AJ, McCauley JW, Russell CA, Smith DJ, Rambaut A. 2014 Integrating influenza antigenic dynamics with molecular evolution. eLife 3, e01914.

[41] Kung HC, Jen KF, Yuan WC, Tien SF, Chu CM. 1978 Influenza in China in 1977: recurrence of influenzavirus A subtype H1N1. Bulletin of the World Health Organization 56, 913–8.

[42] Rozo M, Gronvall GK. 2015 The Reemergent 1977 H1N1 Strain and the Gain-of-Function Debate. mBio 6.

[43] Nakajima K, Desselberger U, Palese P. 1978 Recent human influenza A (H1N1) viruses are closely related genetically to strains isolated in 1950. Nature 274, 334–339.

[44] Park AW, Daly JM, Lewis NS, Smith DJ, Wood JLN, Grenfell BT. 2009 Quantifying the impact of immune escape on transmission dynamics of influenza. Science 326, 726–728.

[45] Wen F, Bedford T, Cobey S. 2016 Explaining the geographical origins of seasonal influenza A (H3N2). Proceedings of the Royal Society B 283, 20161312–9.

[46] WHO. 2014 Influenza Fact Sheet. World Health Organization.

[47] Axelsen JB, Yaari R, Grenfell BT, Stone L. 2014 Multiannual forecasting of seasonal influenza dynamics reveals climatic and evolutionary drivers. Proceedings of the National Academy of Sciences of the United States of America 111, 9538–42.

[48] Du X, King AA, Woods RJ, Pascual M. 2017 Evolution-informed forecasting of seasonal influenza A (H3N2). Science translational medicine 9.

[49] Ranjeva S, Subramanian R, Fang VJ, Leung GM, Ip DKM, Perera RAPM, Peiris JSM, Cowling BJ, Cobey S. 2019 Age-specific differences in the dynamics of protective immunity to influenza. Nature Communications 10, 1660.

[50] Hadfield J, Megill C, Bell SM, Huddleston J, Potter B, Callender C, Sagulenko P, Bedford T, Neher RA. 2017 Nextstrain: real-time tracking of pathogen evolution. bioRxiv p. 224048.

[51] Alonso D, McKane AJ, Pascual M. 2007 Stochastic amplification in epidemics. Journal of The Royal Society Interface 4, 575–582.

[52] Fox JW, Vasseur D, Cotroneo M, Guan L, Simon F. 2017 Population extinctions can increase metapopulation persistence. Nature Ecology & Evolution 1, 1271–1278.

[53] CDC. 2015 Flu Vax View. Centers for Disease Control and Prevention.

[54] Halloran ME, Haber M, Ira M Longini J, Struchiner CJ. 1991 Direct and Indirect Effects in Vaccine Efficacy and Effectiveness. American journal of epidemiology 133, 323–331.

[55] McLean HQ, Thompson MG, Sundaram ME, Meece JK, McClure DL, Friedrich TC, Belongia EA. 2014 Impact of Repeated Vaccination on Vaccine Effectiveness Against Influenza A(H3N2) and B During 8 Seasons. Clinical Infectious Diseases 59, 1375–1385.

[56] Skowronski DM, Chambers C, De Serres G, Sabaiduc S, Winter AL, Dickinson JA, Gubbay JB, Fonseca K, Drews SJ, Charest H, Martineau C, Krajden M, Petric M, Bastien N, Li Y, Smith DJ. 2017 Serial vaccination and the antigenic distance hypothesis: effects on influenza vaccine effectiveness during A(H3N2) epidemics in Canada, 2010-11 to 2014-15. Journal of Infectious Diseases.

[57] Flood EM, Rousculp MD, Ryan KJ, Beusterien KM, Divino VM, Toback SL, Sasane M, Block SL, Hall MC, Mahadevia PJ. 2010 Parents’ decision-making regarding vaccinating their children against influenza: A web-based survey. Clinical therapeutics 32, 1448–1467.

[58] Lewnard J, Cobey S. 2018 Immune History and Influenza Vaccine Effectiveness. Vaccines 6, 28.

[59] Cobey S, Gouma S, Parkhouse K, Chambers BS, Ertl HC, Schmader KE, Halpin RA, Lin X, Stockwell TB, Das SR, Landon E, Tesic V, Youngster I, Pinsky BA, Wentworth DE, Hensley SE, Grad YH. 2012 Poor immunogenicity, not vaccine strain egg adaptation, may explain the low H3N2 influenza vaccine effectiveness in 2012-13. Clinical Infectious Diseases.

[60] Fonville JM, Wilks SH, James SL, Fox A, Ventresca M, Aban M, Xue L, Jones TC, Le NMH, Pham QT, Tran ND, Wong Y, Mosterin A, Katzelnick LC, Labonte D, Le TT, van der Net G, Skepner E, Russell CA, Kaplan TD, Rimmelzwaan GF, Masurel N, de Jong JC, Palache A, Beyer WEP, Le QM, Nguyen TH, Wertheim HFL, Hurt AC, Osterhaus ADME, Barr IG, Fouchier RAM, Horby PW, Smith DJ. 2014 Antibody landscapes after influenza virus infection or vaccination. Science 346, 996–1000.

[61] Chen YQ, Wohlbold TJ, Zheng NY, Huang M, Huang Y, Neu KE, Lee J, Wan H, Rojas KT, Kirkpatrick E, Henry C, Palm AKE, Stamper CT, Lan LYL, Topham DJ, Treanor J, Wrammert J, Ahmed R, Eichelberger MC, Georgiou G, Krammer F, Wilson PC. 2018 Influenza Infection in Humans Induces Broadly Cross-Reactive and Protective Neuraminidase-Reactive Antibodies. Cell 173, 417–429.e10.

[62] Cobey S, Hensley SE. 2017 Immune history and influenza virus susceptibility. Current opinion in virology 22, 105–111.

[63] Zost SJ, Parkhouse K, Gumina ME, Kim K, Diaz Perez S, Wilson PC, Treanor JJ, Sant AJ, Cobey S, Hensley SE. 2017 Contemporary H3N2 influenza viruses have a glycosylation site that alters binding of antibodies elicited by egg-adapted vaccine strains. Proceedings of the National Academy of Sciences.

[64] Linderman SL, Chambers BS, Zost SJ, Parkhouse K, Li Y, Herrmann C, Ellebedy AH, Carter DM, Andrews SF, Zheng NY, Huang M, Huang Y, Strauss D, Shaz BH, Hodinka RL, Reyes-Terán G, Ross TM, Wilson PC, Ahmed R, Bloom JD, Hensley SE. 2014 Potential antigenic explanation for atypical H1N1 infections among middle-aged adults during the 2013-2014 influenza season. Proceedings of the National Academy of Sciences of the United States of America 111, 15798–15803.

[65] Davenport FM, Hennessy AV. 1956 A serologic recapitulation of past experiences with influenza A; antibody response to monovalent vaccine. The Journal of experimental medicine 104, 85–97.

[66] Davenport FM, Hennessy AV. 1957 Predetermination by infection and by vaccination of antibody response to influenza virus vaccines. The Journal of experimental medicine 106, 835–50.

[67] Zarnitsyna VI, Bulusheva I, Handel A, Longini IM, Halloran ME, Antia R. 2018 Intermediate levels of vaccination coverage may minimize seasonal influenza outbreaks. PLOS ONE 13, e0199674.

[68] Smith DJ, Forrest S, Ackley DH, Perelson AS. 1999 Variable efficacy of repeated annual influenza vaccination. Proceedings of the National Academy of Sciences 96, 14001–14006.

[69] McLean HQ, Thompson MG, Sundaram ME, Kieke BA, Gaglani M, Murthy K, Piedra PA, Zimmerman RK, Nowalk MP, Raviotta JM, Jackson ML, Jackson L, Ohmit SE, Petrie JG, Monto AS, Meece JK, Thaker SN, Clippard JR, Spencer SM, Fry AM, Belongia EA. 2015 Influenza vaccine effectiveness in the United States during 2012-2013: variable protection by age and virus type. Journal of Infectious Diseases 211, 1529–1540.

[70] Galvani AP, Reluga TC, Chapman GB. 2007 Long-standing influenza vaccination policy is in accord with individual self-interest but not with the utilitarian optimum. Proceedings of the National Academy of Sciences 104, 5692–5697.

[71] Jackson C, Vynnycky E, Mangtani P. 2010 Estimates of the transmissibility of the 1968 (Hong Kong) influenza pandemic: evidence of increased transmissibility between successive waves. American Journal of Epidemiology 171, 465–478.

[72] Biggerstaff M, Cauchemez S, Reed C, Gambhir M, Finelli L. 2014 Estimates of the reproduction number for seasonal, pandemic, and zoonotic influenza: a systematic review of the literature. BMC Infectious Diseases 14, 1–20.

[73] Carrat F, Vergu E, Ferguson NM, Lemaitre M, Cauchemez S, Leach S, Valleron AJ. 2008 Time lines of infection and disease in human influenza: a review of volunteer challenge studies. American Journal of Epidemiology 167, 775–785.

[74] UN. 2013 World Population Prospects: The 2012 Revision. United Nations, Department of Economic and Social Affairs, Population Division New York.

